# Surround Suppression of Broadband Images

**DOI:** 10.1101/2024.05.15.594329

**Authors:** Victor J. Pokorny, Kimberly B. Weldon, Cheryl A. Olman

## Abstract

Visual perception is profoundly sensitive to context. Surround suppression is a well-known visual context effect in which the firing rate of a neuron is suppressed by stimulation of its extra-classical receptive field. The majority of contrast surround suppression studies exclusively use narrowband, sinusoidal grating stimuli; however, it is unclear whether the results produced by such artificial stimuli generalize to real-world, naturalistic visual experiences. To address this issue, we developed a contrast discrimination paradigm that includes both naturalistic broadband textures and narrowband grating textures. All textures were matched for first order image statistics and overall perceptual salience. We observed surround suppression across broadband textures (*F*(1,6)=19.01, *p*=.005); however, effect sizes were largest for narrowband, sinusoidal gratings (Cohen’s *d*=1.83). Among the three broadband texture types, we observed strongest suppression for the texture with a clear dominant orientation (stratified: Cohen’s *d*=1.29), while the textures with a more even distribution of orientation information produced weaker suppression (fibrous: Cohen’s *d*=0.63; braided: Cohen’s *d*=0.65). We also observed an effect of texture identity on the slope of psychometric functions (*F*(1.98,11.9)=7.29, *p*=0.01), primarily driven by smaller slopes for the texture with the most uniform distribution of orientations. Our results suggest that well-known contextual modulation effects only partially generalize to more ecologically valid stimuli.

## Introduction

Many classic visual illusions, from Adelson’s Checker-Shadow Illusion to the moon illusion, highlight how perception of target stimuli can be profoundly modulated by context. Understanding the mechanisms of context-dependent effects can provide unique insights into the encoding and representation of visual information in the brain.

One well-studied mechanism of contextual modulation is surround suppression. Neurophysiologically, surround suppression is defined as a reduction in firing rate when a neuron’s extra-classical field is stimulated (Cavanaugh et al., 2002). A perceptual analogue of this effect has long been observed in which the percieved contrast of a target stimulus is modulated by the presence of surrounding stimuli. Importantly, laboratory measurements of neural surround suppression in primary visual cortex (V1) generally agree with psychophysical measurements of perceived contrast suppression, suggesting a direct link between perceptual and neural surround suppression (Zipser et al., 1996).

The neural mechanisms of surround suppression can be provisionally categorized into V1-intrinsic and V1-extrinsic mechanisms. Within V1, surround suppression is thought to be mediated by lateral connections between neurons that sample different spatial locations, but have similar orientation tuning preferences (Gilbert & Wiesel, 1983; Smith, 2006). The evidence for such like-to-like inhibition includes a) anatomical evidence of “long-range” connections between neurons within V1 with similar orientation preferences, b) physiological evidence of greater suppression of V1 neural firing for parallel oriented surrounds and c) psychophysical evidence of greater suppression of perceived contrast for parallel surrounds (Shushruth et al., 2013; Smith, 2006). Evidence for V1-extrinsic feedback playing a role in surround suppression is primarily drawn from neurophysiological studies showing that surround suppression effects occur too quickly to be purely mediated by the slow, unmyelinated axons of V1 lateral connections (Bair et al., 2003). Instead, faster, myelinated feedback connections from higher level brain areas could mediate such quick suppression of V1 responses. Furthermore, higher level stimulus features (e.g., segmentation cues) and attentional manipulations are known to impact the degree of contextual modulation in V1 (Coen-Cagli et al., 2015; Flevaris & Murray, 2015; Reynolds & Heeger, 2009).

Most studies of contextual modulation have used artificial, narrowband stimuli. These stimuli are powerful because the experimenter can tightly control the physical characteristics of the stimuli (e.g., spatial frequency, orientation, phase, size, etc.); however, the extent to which findings derived from grating stimuli generalize to more naturalistic and complex stimuli is unclear. Sinusoidal gratings are rarely encountered outside of the psychophysics laboratory and are likely to engage extrastriate surround suppression mechanisms differently than naturalistic textures. In particular, there is growing evidence that V2 neurons show some degree of texture selectivity such that comparing perceived surround suppression across different texture types may allow for inferences regarding suppression in areas beyond V1 (Ziemba et al., 2016). For these reasons, we developed a behavioral task that uses both naturalistic, broadband textures and narrowband sinusoidal gratings. The task allowed for manipulation of surround orientation (parallel vs. orthogonal) to assess the effect of broadband and sinusoidal textures on both orientation-tuned and untuned surround suppression. Additionally, to quantify how higher order image statistics impact the degree of suppression, we included phase-scrambled (noise) control conditions (Herrera-Esposito et al., 2021; Julesz, 1981). We predicted surround suppression would be stronger for conditions in which visual stimuli have surrounds that better predict the center (i.e., parallel orientation) across both broad and narrowband stimuli. Furthermore, we predicted that higher order image statistics would be important for robust surround suppression.

## Methods

### Participants

Twelve participants (6 male, 6 female; mean age: 29 years) with normal or corrected to normal vision participated in the study. Four participants were the authors and familiar with participating in psychophysical tasks. Non-author participants were naïve to the purpose of the experiment. Seven participants completed all 18 conditions while the remaining five completed only a subset of the conditions. Participants provided written informed consent prior to participation and were compensated $15/hour. All procedures were approved by the University of Minnesota Institutional Review Board and are in accordance with ethical standards of the Declaration of Helsinki.

### Apparatus

Stimuli were displayed on a 61 cm LCD monitor (EIZO FlexScan SX2462W) with a resolution of 1920x1280 and refresh rate of 60Hz, controlled by a 21 inch iMac with an NVIDIA GeForce GT650M graphics card with 8 bit DVI output. A Bits# processor was used to provide 10-bit resolution control of contrast level. Mean luminance of the screen was 72 cd/m^2^. The monitor was calibrated using an optical photometer (Photo Research SpectraScan 655). Viewing distance was 70 cm.

### Texture Selection Process

Images were downloaded in July 2018 from the Describable Texture Database (Cimpoi et al., 2013), a growing repository of textures categorized by the words that human observers use to describe them. There were 5,650 images divided into 48 categories (e.g., ‘braided’, ‘porous’, and ‘bubbly’). Using custom Python code, we quantified the following statistics on the pixel samples: spatial variance of image contrast, peak spatial frequency, proportion of low spatial frequencies, distribution of Fourier components across orientation, and the skew, kurtosis, and percentage of voxels clipped by scaling (more information on these statistics can be found in the Supplemental Material). We selected texture images that met the following criteria: reasonably uniform contrast across image (defined as a spatial variance of < 2), pixel histogram with low skew and kurtosis (-0.5 < skew < 0.5, -1 < kurtosis < 1), peak spatial frequency between 12 and 16 cycles per image, >= 25% of power near peak spatial frequency, < 25% of power in DC. The qualifying textures used in the final experiment were ‘braided’, ‘stratified’, and ‘fibrous’ textures. These stimuli represented a spectrum of ‘oriented’ to ‘non-oriented’ textures, with stratified textures as the most oriented, fibrous as the least oriented and braided as intermediate (Figure 1). Note that the terms ‘oriented’ and ‘non-oriented’ refer to how evenly-distributed the Fourier magnitudes were across orientations, with non-oriented textures having more uniformly distributed magnitudes.

**Figure 1.**
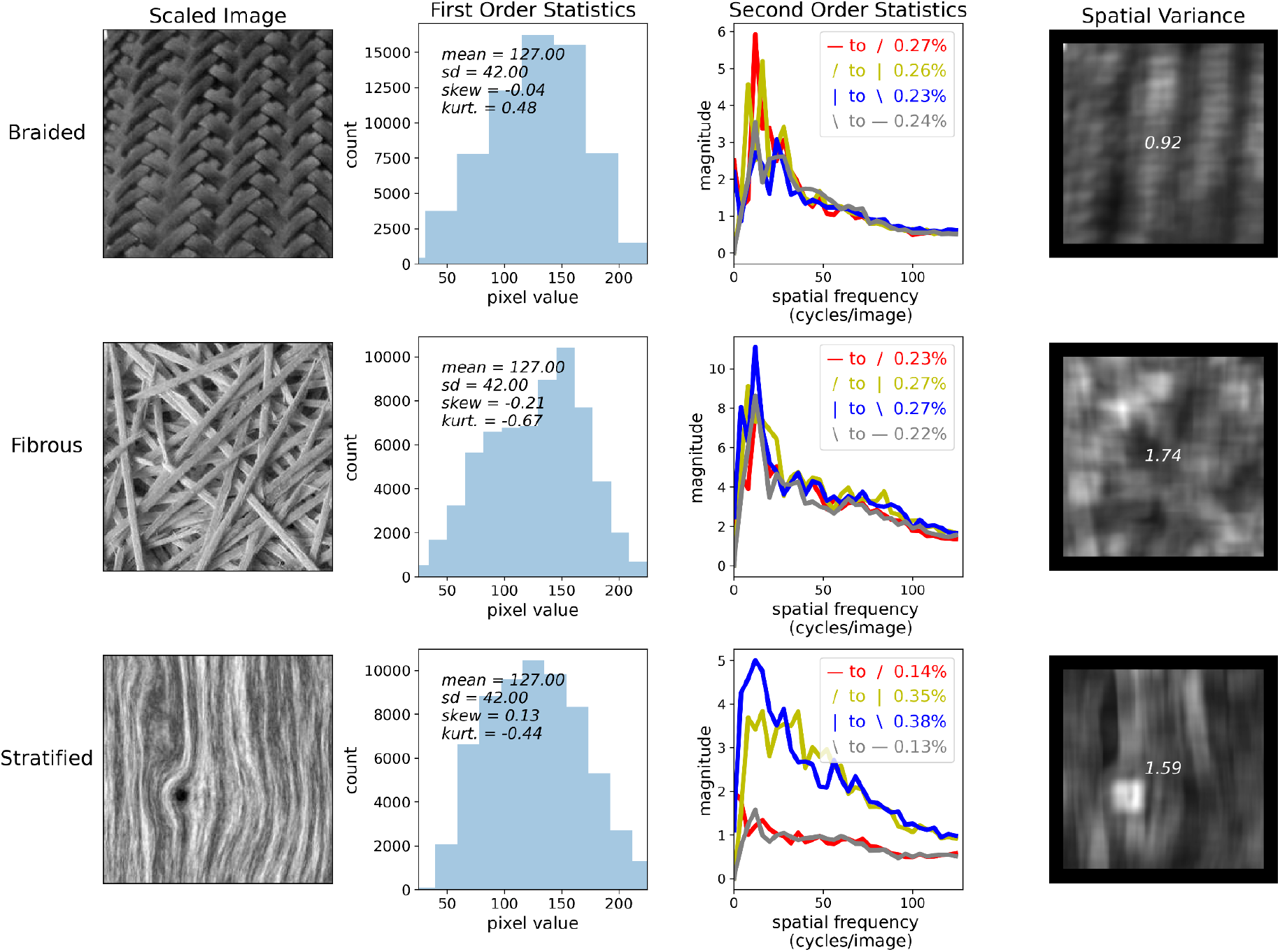
Selection of Texture Stimuli. Scaled images of the textures selected from the Describable Texture Database (DTD) are represented in the first column. Histograms in column two depict the pixel intensity distributions of each image. Column three depicts the fourier magnitudes for each image averaged within four orientation bins (red: 0 < *θ* ≤ π/4, yellow: π/4 < *θ* ≤ π/2, blue: π/2 < *θ* ≤ 3π/4, gray: 3π/4 < *θ* ≤ π). These bins are represented in the legends as ‘–’ representing 0 radians, ‘/’ representing π/4 radians, ‘|’ representing π/2 radians and ‘\’ representing 3π/4 radians. The percentage in the legend is the integral of the magnitude spectrum within a given orientation bin relative to the total integrated magnitude spectrum of the image. The braided and fibrous textures contained Fourier magnitudes that were roughly equivalent across bins while the stratified texture contained larger Fourier magnitudes in the vertical direction. In the fourth column, each pixel represents the variance of a 32 x 32 pixel sliding window (pixels close to the edge are set to zero because the window extends outside the bounds of the image). Spatial Var. is the total variance of the individual sliding window variances for a given image; lower values indicate greater uniformity of contrast across the image. These textures were selected, in part, because they had relatively low spatial variance.

### Stimulus Creation Process

A 300x300 pixel sinusoidal grating was created with random orientation and phase and a frequency of 14 cycles/image. Ten 256x256 pixel exemplars were created from the original 300x300 pixel image by randomly selecting a 256x256 region of the original image. Each exemplar was ‘Gaussianized’ by ranking the pixel intensities and replacing them by pixels drawn from a (similarly ranked) normal distribution. The resulting ‘grating’ stimuli had a peak spatial frequency of 13.6 cycles/256 pixels (i.e., centered in the same 12–16 cycles/image band) and some power at higher frequencies because the Gaussianization resulted in gratings that no longer had perfect sinusoidal luminance modulation (Figure 2).

**Figure 2.**
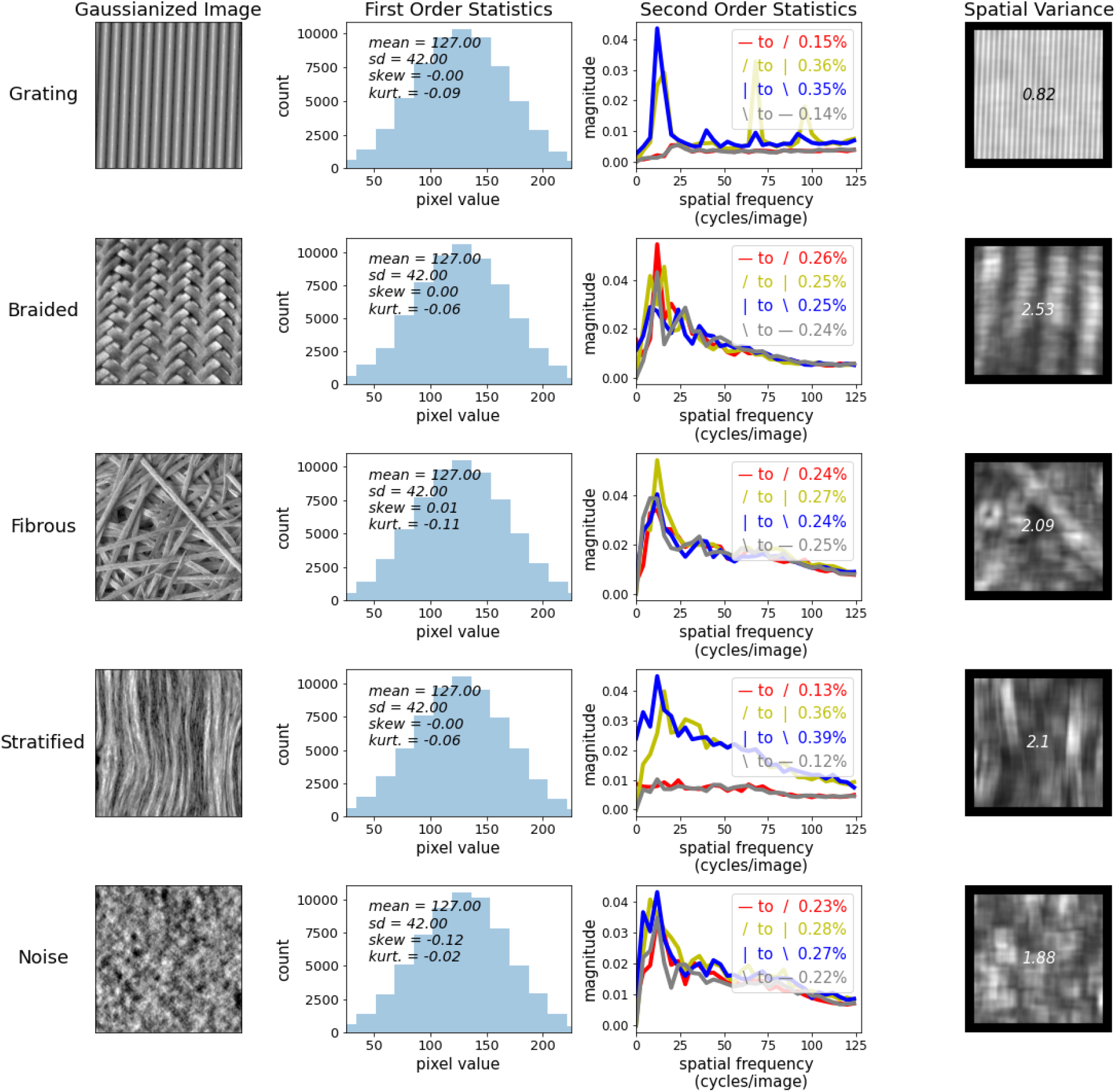
Image Statistics of Gaussianized Task Stimuli. The first column shows the Gaussianized versions of the stimuli that were used for the task. The second column shows the Gaussianized pixel distributions and corresponding moments for each of the textures. The third column shows the fourier amplitude spectra of the stimuli. Note that Gaussianizing the grating stimuli introduced harmonic magnitude spikes. The fourth column shows the spatial variance of the textures. In general, the spatial variance of the textures increased due to the Gaussianization process (as can be seen by comparing values of Figure 1 with Figure 2).

Texture samples were subjected to the same process of selecting random 256x256 windows from 300x300 pixel images downloaded from the Describable Texture Database. These samples were also Gaussianized by replacing the pixel intensity values with normally distributed pixel intensities. Thus, the first order statistics were roughly matched across textures (Figure 2). A noise sample was created for each texture by taking the Fourier transform of the texture, replacing the phase portion of the spectrum with random values uniformly distributed between -π and π (taking care to respect the Hermitian symmetry of the matrix).

### Stimulus presentation

256x256 pixel images were presented using PsychoPy v1.85.2 and were scaled to subtend 5 degrees of visual angle. Thus, spatial frequencies of 12–16 cycles/image corresponded to spatial frequencies of 2.4–3.2 cycles/degree of visual angle. All stimuli consisted of a central circular texture with or without an annular surrounding texture. Each center had a 2 degree diameter and each surround had a 5 degree diameter. There was a gap of 0.25 degrees between center and surround, created by a raised cosine mask. Black circles delineated the center regions throughout the experiment. The black circles likely act as segmentation cues between center and surround which are known to diminish degree of suppression; however, we included the black circles to reduce the influence of figure-ground modulation. In parallel conditions, center and surround had the same orientation. For orthogonal conditions, the orientation of the surround was 90 degrees offset from the center. Eighteen stimulus conditions were used in this study (see Panel B of Figure 3).

**Figure 3.**
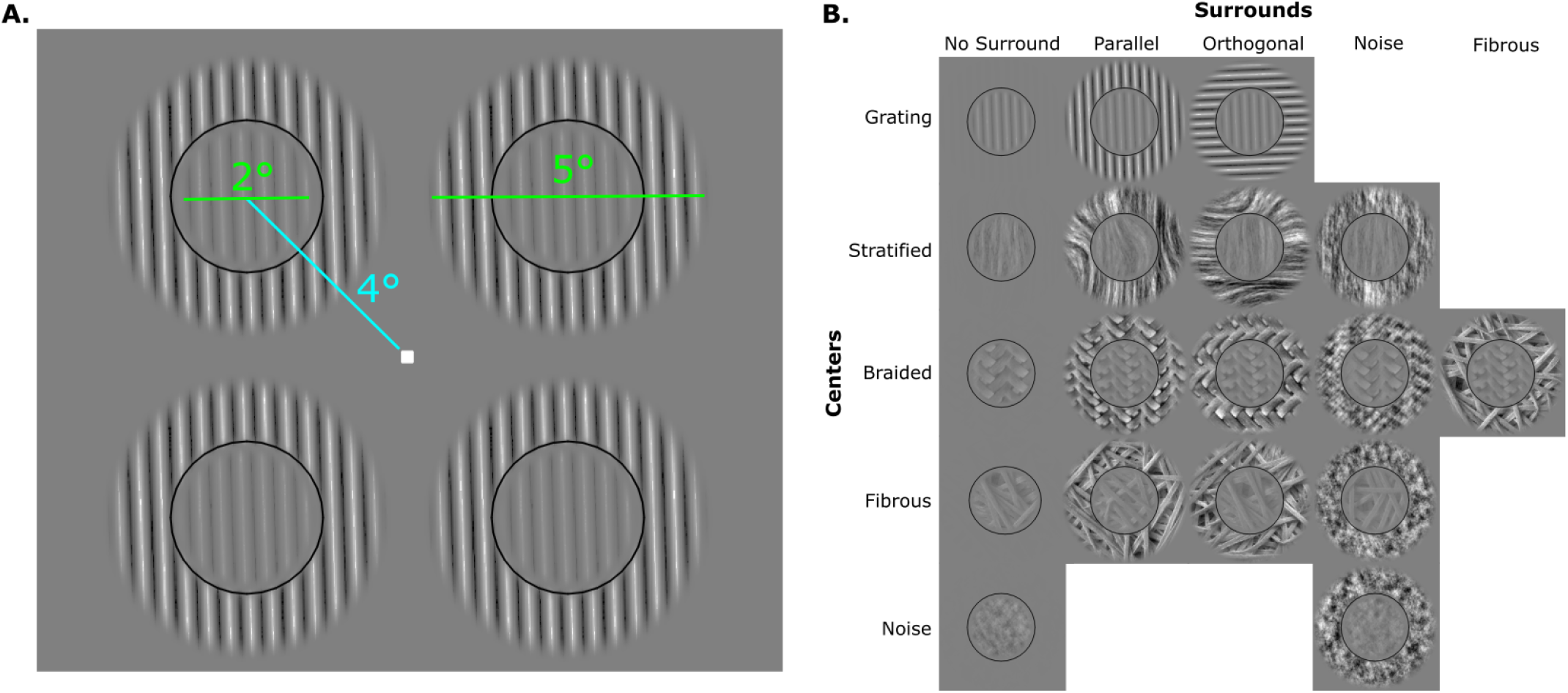
Task Examples. Panel A shows the stimulus array for an example condition (grating; center with parallel surround). Teal line indicates that the stimuli were presented at 4° eccentricity. The diameter of the centers subtended 2° of visual angle and the diameter of the surrounds subtended 5°. Panel B shows all the conditions presented for the task.

### Procedure

A four-alternative forced choice (4AFC) paradigm was used to determine contrast discrimination thresholds (Jäkel & Wichmann, 2006; Vancleef et al., 2018). During each trial, four stimuli were simultaneously presented for a duration of 300 ms at 3 degrees eccentricity from a central white fixation point (Figure 3). One target stimulus had higher RMS contrast relative to the three distractors. The RMS contrast for each stimulus was calculated as the standard deviation of the pixel intensity distribution divided by 128 (the maximum deviation from the mean pixel value of 127). The degree to which the target’s contrast differed from the distractors’ contrast was referred to as the delta contrast (dC). In each trial, the position of the target was randomized. For conditions with a surround, the surround contrast was constant, at 0.32 RMS contrast.

The participant was asked to indicate which stimulus had higher contrast compared to the three distractors by pressing a corresponding keyboard key. If the response was correct, the fixation point briefly changed to black. An incorrect response resulted in no change to the fixation point. The first run, consisting of 38–45 trials, used a one-up-one-down staircase to narrow the range of dCs necessary to target the participants’ estimated threshold. This initial run determined a set of 11 log-spaced dCs, centered on the initial threshold estimate to be measured in the subsequent three runs for each participant at each pedestal contrast level for each condition. These subsequent runs consisted of 75 trials each. This procedure was used to ensure uniform sampling across the psychometric function to allow estimation of slope as well as threshold. We did not use the staircase handlers built into PsychoPy because we could not know ahead of time whether the settings designed for gratings would work similarly for textures. Each participant completed at least four runs of the task. Seven participants completed each condition, averaging 286 (240–375) trials (distributed across 11 dC values) for each of nine pedestal contrast levels (0, 0.01, 0.02, 0.03, 0.04, 0.06, 0.08, 0.12, and 0.16).

### Threshold and Slope Estimation

We fit a logistic function (percent correct vs. dC (Figure 4)) for each observer, condition and pedestal. Thresholds were defined at 62.5% accuracy which is halfway between chance (25%) and perfect performance (100%). Slopes of the logistic functions were clamped at ten to ensure proper fitting. Excluding cases where slopes were clamped did not substantially change our results.

**Figure 4.**
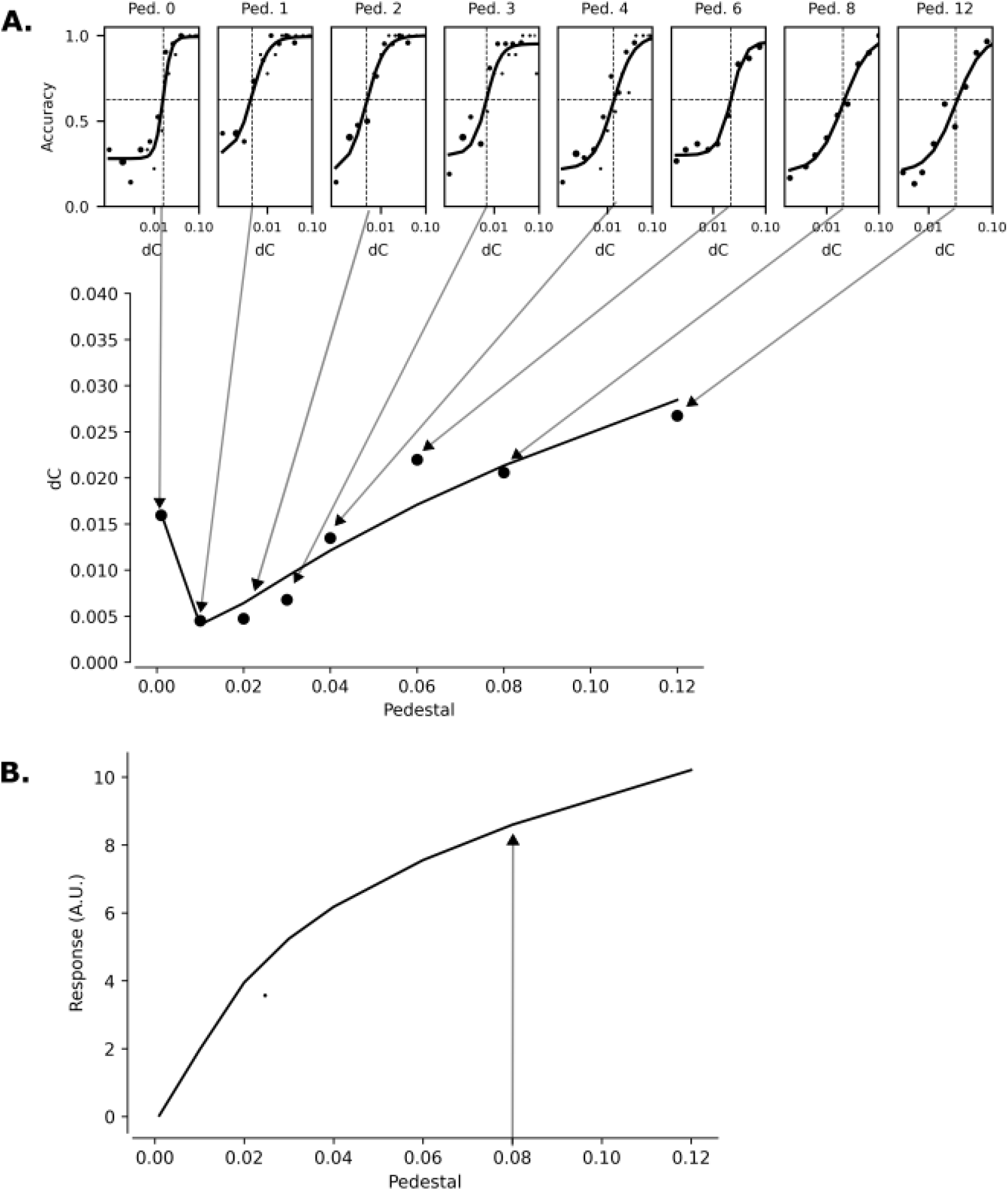
Modeling of Logistic Functions and TvC curves. Panel A depicts a fitted logistic function for each pedestal for a single subject and condition. Below the logistic functions is a TvC curve fitted to the thresholds from the logistic functions. Panel B depicts the Naka-Rushton function which is the integral of the reciprocal of the TvC curve in Panel A. To make condition comparisons we extracted the response value at 8% contrast (denoted by the arrow in Panel B) for each individual.

### Response Estimates

The Naka-Rushton formula (see figure 4), *R* = (*A* × *C*^*p*+*q*^)/(*s*^*q*^ + *C*^*q*^), is commonly used to represent neural and behavioral responses as a function of contrast because it captures both accelerating non-linearity at low contrast and saturating (i.e., decelerating) non-linearity at high contrast (see Panel B of Figure 4). The reciprocal of the derivative of this four-parameter function describes the relationship between behavioral thresholds and pedestal contrast (see Panel A in Figure 4). Hereafter we refer to this function as a threshold vs. contrast (TvC) curve. The TvC curve was fit to the discrimination thresholds at the lowest 8 pedestals (0 - 0.12 contrast) to avoid the region of the contrast-response function where suppression is expected to change into facilitation as the contrast of the central target approaches the contrast of the surround. Functions were fitted using Python scipy.optimize.curve_fit, which uses non-linear least squares to estimate fit parameters. For the four parameter TvC curve, the ranges used for the parameters were *A* (0, 50), *p* (0.05, 1.0), *q* (0.50, 2.5), and *s* (0, 0.10). We then used the estimated response at 8% RMS contrast as the dependent variable for statistical comparison.

### Statistical Comparisons

Statistical comparisons between conditions were performed using R-4.0.5 (R Core Team, 2023). Omnibus repeated measures ANOVAs were computed using the aov_ez function from the afex package with texture and orientation as within subject factors (Singmann et al., 2023). We computed follow-up pairwise t-tests using the base R t.test function. For pairwise comparisons, we also computed Cohen’s *d* as the mean of the difference scores between two conditions divided by the standard deviation of the difference scores: mean(X_condition1_-X_condition2_) / sd(X_condition1_-X_condition2_). Finally, we computed Hedges’ *g* using the hedges_g function from the esc package (Lüdecke, 2019). This function takes the Cohen’s d value as input and applies a small sample size bias correction in which *d* is multiplied by 1 - (3 / (4 * *n* - 9)) where *n* is the number of subjects.

### Inclusion criteria

Only a small number of fits required one of the 4 TvC parameters set to the limit of the allowed range (A: 1 of 126, p: 6 of 126, q: 2 of 126, *s*: 1 of 126). Excluding these cases from the group averages did not substantially change our results. Similarly, excluding the few individual discrimination thresholds for which the logistic fit had high error did not impact the results reported below.

## Results

### Thresholds

The selected textures were roughly equivalent in terms of perceptual salience: contrast discrimination thresholds for no-surround stimuli did not differ as a function of narrowband, broadband or noise conditions (see Figure 5; *F*(1.82,10.93)=0.48, *p*=0.61). Furthermore, the relationship between contrast discrimination threshold and pedestal contrast was described by similar *p* and *q* parameters in the Naka-Rushton formula used to estimate the magnitude of the visual response evoked by each texture/surround condition as a function of contrast (see Supplemental Figure 2).

**Figure 5.**
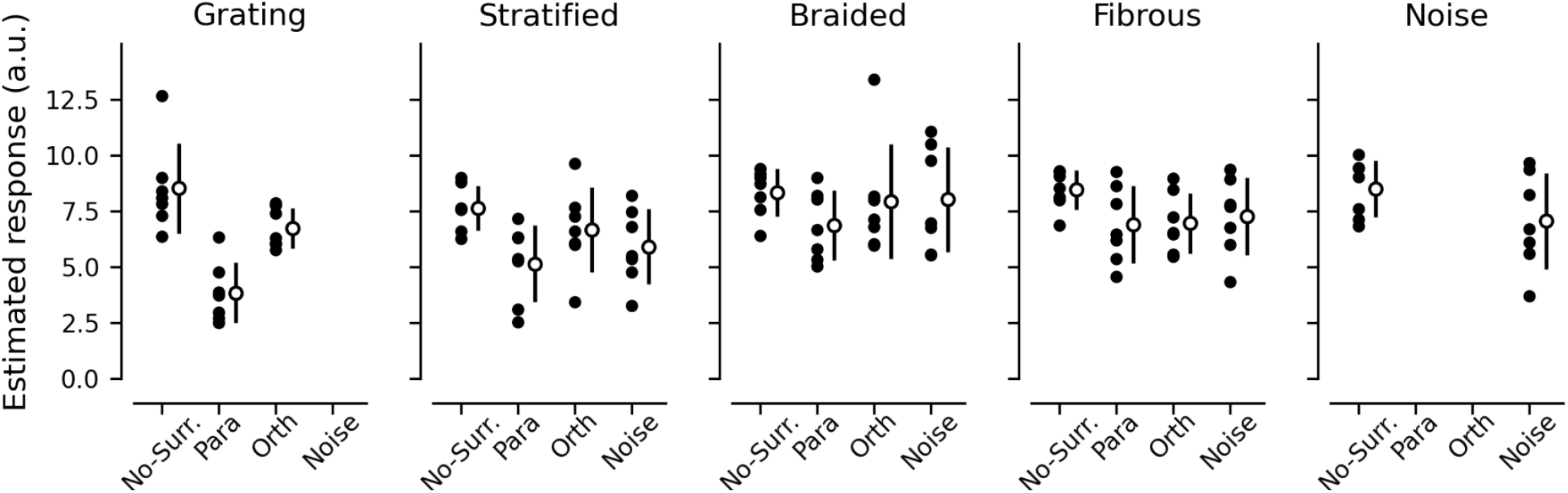
Estimated Responses for All Conditions. Black dots represent estimated responses to an 8% RMS contrast stimulus for each individual for a given condition. White dots and black lines represent mean and standard deviation, respectively. No-Surr: No-surround, Para: Parallel, Orth: Orthogonal.

For the grating stimuli, we successfully replicated the canonical surround suppression effect: decreased perceived contrast of an 8% RMS contrast target when the center was accompanied by a parallel surround (*t*(6)=4.85, *p*=<.001, *d*=1.83, *g =* 1.54). Similarly, we observed an expected decrease in perceived contrast for parallel surrounds relative to orthogonal surrounds (*t*(6)=5.64, *p*=<.001, *d* = 2.13, *g =* 1.79).

For broadband stimuli (i.e., stratified, braided, and fibrous textures), we observed a reduction in perceived contrast for the parallel condition relative to no-surround (*F*(1,6)=19.01, *p*=.005). Of the three broadband textures, the stratified stimuli led to the strongest suppression (*t*(6)=3.41, *p*=0.01, *d* = 1.29, *g* =1.09) while the braided and fibrous textures produced roughly equivalent suppression (braided: *t*(6)=1.73, *p*=0.07, *d* = 0.65, *g* = 0.55; fibrous: *t*(6)=1.67, *p*=0.07, *d* = 0.63, *g* = 0.53). We observed only a trend for broadband stimuli producing orientation-dependent suppression (i.e., parallel vs. orthogonal contrast; *F*(1,6)=3.84, *p*=0.098) with stratified textures again producing the largest effect sizes (stratified: *t*(6)=1.64, *p*=0.08, *d*=0.62, *g* = 0.52; braided: *t*(6)=0.88, *p*=0.21, *d*=0.33, *g* = 0.28; fibrous: *t*(6)= -0.01, *p*=0.5, *d* = 0, *g* = 0).

Because phase scrambling a texture can make it unrecognizable, we hypothesized that phase scrambled surrounds would be ineffective suppressors for their unscrambled (intact texture) counterparts. In line with this hypothesis, broadband texture thresholds were not significantly suppressed by the phase scrambled surround conditions relative to no-surround conditions (F(1,6)=2.9, p=0.14). However, we did not observe a significant difference between phase-scrambled surrounds and intact parallel surrounds (*F*(1,6)=2.94, *p*=0.14). This suggests that phase-scrambled surrounds created intermediate suppression: not strong enough to significantly differ from no-surrounds, but not weak enough to significantly differ from parallel surrounds. Finally, we did not observe a significant suppressive effect when both the center and surround were phase scrambled (noise-noise *t*(6)=1.3, *p*=0.12, *d*=0.49, *g* = 0.41). In each of these comparisons, the mean responses trended in the expected direction such that it is possible that larger sample sizes would reveal the hypothesized effects.

### Slopes

In addition to perceptual threshold estimates, we also assessed the degree to which slope estimates of the fitted logistic functions differed across narrowband and broadband textures (Figure 6). We observed a significant difference in slopes across all textures (*F*(2.4,14.4)=5.3, *p*=0.02). Individual t-tests revealed significant differences between the grating and fibrous conditions (*t*(6)=2.99, *p*=0.02, *d* = 1.13, *g* = 0.95) and the grating and noise conditions (*t*(6)=3.07, *p* = 0.02, *d* = 1.16, *g* = 0.98). Thus, the fibrous and noise conditions had lower slopes than the grating conditions which is suggestive of less precision in perceptual judgements for fibrous textures. It is worth noting, however, that variance between observers was also lower in the fibrous and noise conditions, suggesting that individual variability in perceptual decision making might be contributing to the higher slope estimates for the more strongly oriented textures.

**Figure 6.**
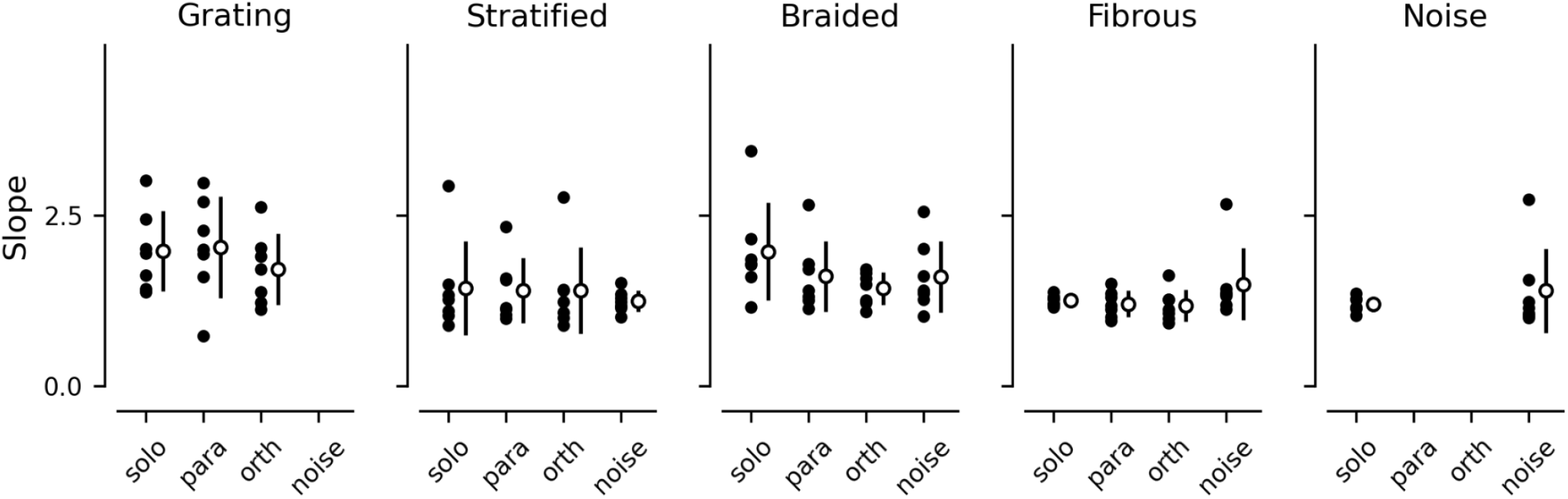
Slopes of Psychometric Functions for All Conditions. Black dots depict the slope of the psychometric function for each individual for a given condition (averaged across contrast pedestals).

## Discussion

Artificial, narrowband gratings allow for fine-grained control of stimulus features known to be encoded by V1 (e.g., orientation, contrast, phase, spatial frequency, size, etc.) and thus are useful for maximizing internal validity of a given experiment. However, experiments using such gratings suffer from poor external validity because the stimuli are rarely encountered outside of the psychophysics laboratory. In contrast, there is a wealth of recent work, primarily inspired by the success of deep neural networks such as Alexnet and VGG16, that attempt to characterize visual perception of “natural scenes” (Allen et al., 2022; Krizhevsky et al., 2017; Roth et al., 2022; Roth & Merriam, 2023; Simonyan & Zisserman, 2014). These natural scene paradigms have superior external validity, but poorer internal validity because the mechanisms by which neural networks accomplish visual tasks and the degree to which these mechanisms extend to the brain are unclear. Examinations using natural textures and other naturalistic stimuli serve as an important bridge between grating and natural scene studies of visual perception.

We created naturalistic broadband texture stimuli with similar statistical properties to narrowband sinusoidal gratings. By selecting textures based on a variety of image statistical criteria (pixel uniformity, skew, kurtosis, peak spatial frequency, etc.) and explicitly matching first order statistics, we ensured internally valid comparisons across narrowband and broadband textures. Moreover, we found that estimated responses for the no-surround narrowband and broadband stimuli did not significantly differ from each other, suggesting roughly equivalent perceptual salience of the stimuli.

Naturalistic, broadband textures induced a classical surround suppression effect; however, the magnitude of suppression was weaker for broadband stimuli (Cohen’s *d*s = .63-1.29) relative to narrowband stimuli (Cohen’s *d* = 1.83). Within the broadband textures, the stratified texture produced a surround suppression effect size roughly twice that of the fibrous and braided textures. Importantly, this stratified texture was the most ‘oriented’ (i.e., had the most uneven distribution of orientation magnitudes giving the texture a clear dominant orientation). In contrast, the textures that produced the weakest suppression (braided and fibrous) had a roughly equal distribution of magnitude across orientations. One likely explanation for this pattern of results is that inhibition from like-to-like lateral connections within V1 was dampened for ‘unoriented’ textures, leading to weaker suppression of perceived contrast. Another way of framing this finding is that narrowband gratings may exaggerate surround suppression effects while more naturalistic textures and visual scenes tend to produce substantially less suppression than would be expected based on narrowband grating studies.

Phase scrambling the surround did not lead to significant changes in the degree of surround suppression relative to intact surrounds. Recent work in both primate and human models has suggested that V2 neurons are more selective for higher order naturalistic texture statistics relative to V1 neurons. In particular, Ziemba et al. (2016) showed that V1 firing rates remain generally constant for both naturalistic textures and phase scrambled noise textures while V2 neurons exhibited increased firing rates for the naturalistic texture as compared to noise. One might expect such modulation of V2 firing rates to impact downstream contrast perception; however, this was not corroborated by our findings. Encouragingly, our results align with recent work by Herrera-Esposito et al. (2021) in which scrambling higher order statistics of textures also did not lead to strong changes in the degree of suppression; however, the authors noted that the effect of phase scrambling seemed to be quite variable from texture to texture. Further work is needed to determine whether manipulating higher order statistics of textures produces meaningful perceptual changes in suppression.

We did not observe strong suppression of perceived contrast when a noise center was surrounded by a noise surround. This result is in disagreement with prior studies (Barch et al., 2012; Dakin et al., 2005). However, these prior studies had larger sample sizes such that it is possible that we were underpowered to detect a true effect. Furthermore, the present task was a 4AFC contrast discrimination task while the Dakin et al. (2005) and Barch et al. (2013) tasks were 2 interval forced choice contrast judgment tasks. Thus, our stimulus configuration may have required a broader deployment of spatial attention which is known to affect surround suppression (Reynolds & Heeger, 2009). Additionally, we presented stimuli peripherally while both Dakin et al. (2005) and Barch et al. (2013) presented stimuli foveally. Previous findings suggest that surround suppression of contrast is stronger in the periphery and is less orientation-specific relative to suppression in the fovea (Xing & Heeger, 2000). Based on these findings, we would expect *stronger* suppression for the present study. The size of the surround is also known to moderate suppression strength (Xing & Heeger, 2001). In the present study, the center diameter was roughly 40% of the surround diameter while for Dakin et al. (2005) the center diameter was roughly 16% of the surround diameter. Thus, the smaller expanse of the surround relative to the center in the present study may have led to weaker suppressive effects.

Unexpectedly, we found that the slopes of the psychometric functions were lower for the fibrous and noise conditions relative to grating stimuli. We are not aware of any reports showing this effect. Less steep slopes in the fibrous and noise conditions suggests that participants were less reliable in their ability to detect contrast increments in those conditions. It is possible that certain texture properties led to more or less reliable perception of contrast. In the case of the fibrous condition, the texture was composed of a complex array of partially occluded cross-oriented edges that may have invoked competing or disrupting mechanisms such as contour integration and/or object recognition. It is also possible that more visually complex textures require more processing time to successfully compute contrast. Further work might investigate how stimulus presentation durations affect the slopes of contrast detection functions for naturalistic textures.

### Future Directions

Although the present study measured psychophysical thresholds, neurophysiological responses (e.g., M/EEG, fMRI, etc.) to broadband stimuli may also prove valuable for clarifying the neural mechanisms of contextual modulation. For example, these stimuli can be used to generate encoding models of the visual cortex which can then be falsified by fMRI or EEG responses (Freeman et al., 2013). Furthermore, a variety of mental health disorders are thought to be associated with altered contextual modulation of contrast (Dakin et al., 2005; Pokorny et al., 2023; Salmela et al., 2021), but none of these reports have used naturalistic stimuli. This gap in the literature is surprising because stimuli that more closely resemble ‘real-world’ visual experiences may better predict real-world functioning of people with mental health disorders. Thus, the study of contextual modulation of broadband textures holds great promise for both basic neuroscientific and applied clinical research.

## Acknowledgements

This work was supported by the following grants from the National Institutes of Health: R01MH112583 and R01NS123482. We would like to thank Emily S. Semaya for early work on this manuscript.

