## Supplemental Materials for "Surround Suppression of Broadband Images"

### *Quantification of Texture Statistics*

**Spatial variance of image contrast**. Images were scaled by subtracting the mean pixel intensity value and dividing by the standard deviation. A map of variance over the central 212x212 pixels of the image sample was then created by roving a 32x32 pixel square over the image, and writing the variance of the (scaled) pixel intensities at the center of each window location. This variance map was divided by the maximum value, and then a single 'contrast variance' index was created by multiplying 100 by the variance of the variance map. Visual inspection guided the selection of 2.0 as a threshold under which textures were similar across the entire 256x256 sample.

**Peak spatial frequency**. Images were 2D Fourier transformed, and the Fourier spectrum was computed by taking the magnitude of the Fourier transform and averaging across orientation. Average power was computed in bands that were 4 cycles per image wide spanning 0 to 128 cycles/image (i.e., the corners of the 2D Fourier spectrum were not considered). The peak spatial frequency was defined as the band with the largest amplitude (collapsed across orientation). Images with a peak spatial frequency band of 12-16 cycles/image were selected for the experiment.

**Peakiness of distribution**. We wanted to select images with relatively narrow spatial frequency distribution, so we created a metric for how concentrated the Fourier spectrum was around the peak spatial frequency. This 'peakiness' metric was computed as the ratio of the sum of the Fourier amplitude spectrum in 5 bins (bin size = 4 cycles/image) centered on the peak spatial frequency to the sum of the entire amplitude spectrum. For example, for an image with a peak spatial frequency in the 12-16 cpi band, the sum of the 5 values representing the amplitudes from 4-24 cpi was divided by the sum of all amplitudes. Images with at least 25% of the Fourier amplitude spectrum contained in the 4-24 cpi band were considered candidates for the experiment.

**Proportion of low spatial frequencies**. We wanted to exclude images with strong gradients or texture discontinuities across the image. Therefore, we computed a metric to exclude images with a lot of power in very low spatial frequencies. To do this, we took the ratio of the amplitude in the lowest spatial frequency bin (0-4 cycles/image) to the amplitude in the peak spatial frequency bin. Images with a low:peak ratio greater than 25% were excluded from the experiment.

**Distribution of Fourier components across orientation**. The 2D Fourier transform was divided into four quadrants (because of symmetry in the Fourier domain, this was actually 4 "propellers" that were symmetric across the center of the Fourier spectrum and divided each half into 4 parts). After computing polar angle as the inverse tangent of x/y, the polar angle map ($\theta$) was divided into ranges from $0 < \theta \leq \pi/4$, $\pi/4 < \theta \leq \pi/2$, $\pi/2 < \theta \leq 3\pi/4$, $3\pi/4 < \theta \leq \pi$. Within each of these (symmetric) masks, the magnitude of the Fourier spectrum was averaged, and then the percentage of the magnitude contained within each band was reported. Note that these bands were not centered on vertical or horizontal. This makes for some non-intuitive mapping between perception of the image samples and the reporting of the Fourier magnitude in each band, but remains a valid metric of orientation homogeneity. Because images were randomly oriented on each trial, the exact definition of vertical is irrelevant. This metric was computed for each image but not used to select candidates for the experiment, because we wanted to measure the effect of orientation distribution on perception.

**Skew, kurtosis, and percentage of voxels clipped by scaling**. Even though the pixel intensity distribution of each image was going to be normalized before presentation, we wanted to select images that would not be strongly altered by this manipulation. We therefore computed the skew and kurtosis of the pixel intensity distribution, as well as the percentage of pixels that would be clipped (set to 0 or 255) when the (non-normalized) image was presented at 33% RMS contrast (i.e., 127 + 42*(img - mean(image))/stdev(img)). Images with -0.5 < skew < 0.5, -1. < kurtosis < 1, and no more than 1% of the voxels clipped by scaling were considered candidates for the experiment.

*Texture Selection*

29 candidate textures were selected based on image statistics (see Supplemental Figure 1). We then selected 3 of these 29 that appeared to be perceptually uniform. One (fibrous_0165.jpg) was selected because it appeared non-oriented and the computed orientation distribution was uniform. One (braided_0102.jpg) was selected because it appeared oriented even though the computed orientation distribution was uniform. The third (stratified_0128.jpg) was selected because it appeared oriented and the Fourier analysis confirmed that vertical orientations were strongly represented, compared to horizontal.

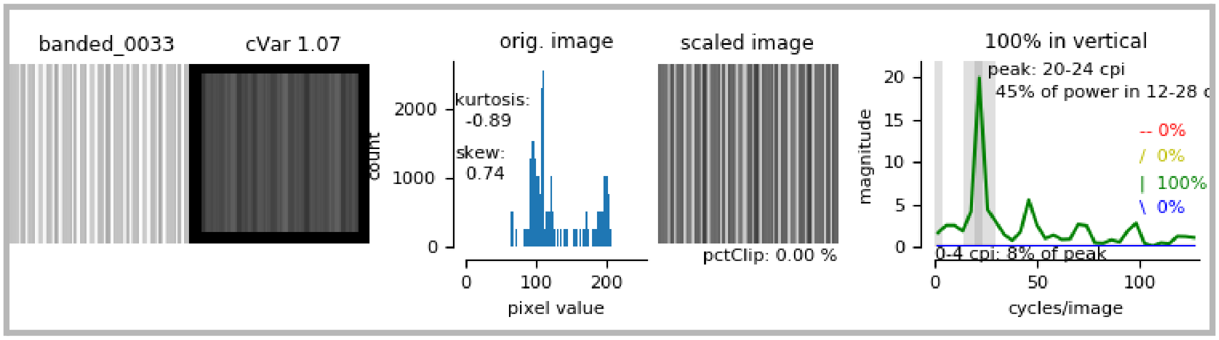
banded_0033
cVar 1.07
orig. image
kurtosis: -0.89
skew: 0.74
count
pixel value
scaled image
pctClip: 0.00 %
100% in vertical
peak: 20-24 cpi
45% of power in 12-28 c
-- 0%
/ 0%
| 100%
\ 0%
0-4 cpi: 8% of peak
magnitude
cycles/image

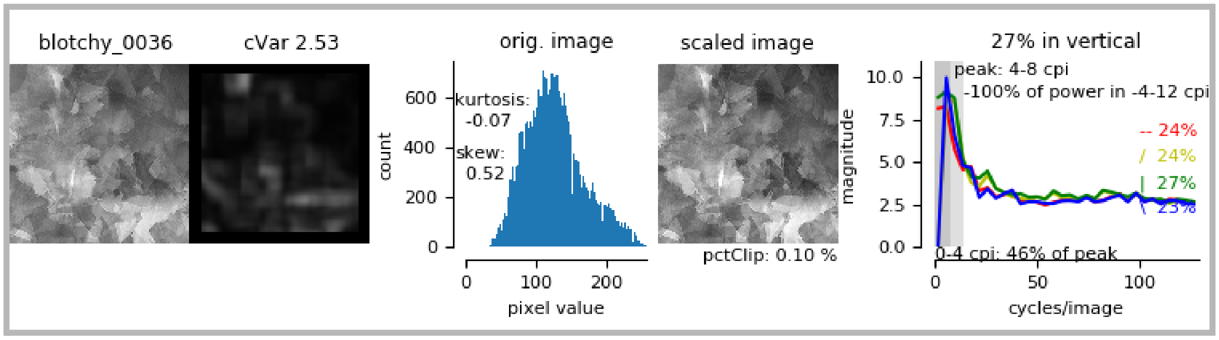
blotchy_0036
cVar 2.53
orig. image
kurtosis: -0.07
skew: 0.52
count
pixel value
scaled image
pctClip: 0.10 %
27% in vertical
peak: 4-8 cpi
100% of power in -4-12 cpi
-- 24%
/ 24%
| 27%
\ 23%
0-4 cpi: 46% of peak
magnitude
cycles/image

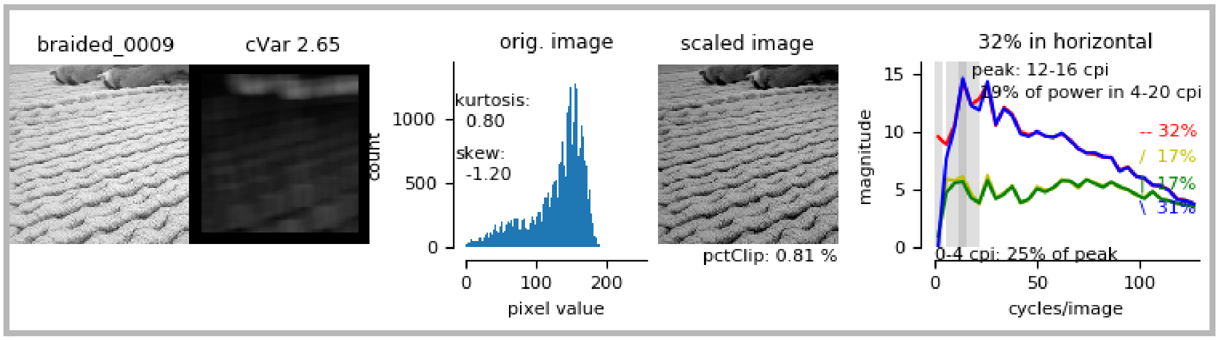
braided_0009
cVar 2.65
orig. image
kurtosis: 0.80
skew: -1.20
count
pixel value
scaled image
pctClip: 0.81 %
32% in horizontal
peak: 12-16 cpi
19% of power in 4-20 cpi
-- 32%
/ 17%
| 17%
\ 31%
0-4 cpi: 25% of peak
magnitude
cycles/image

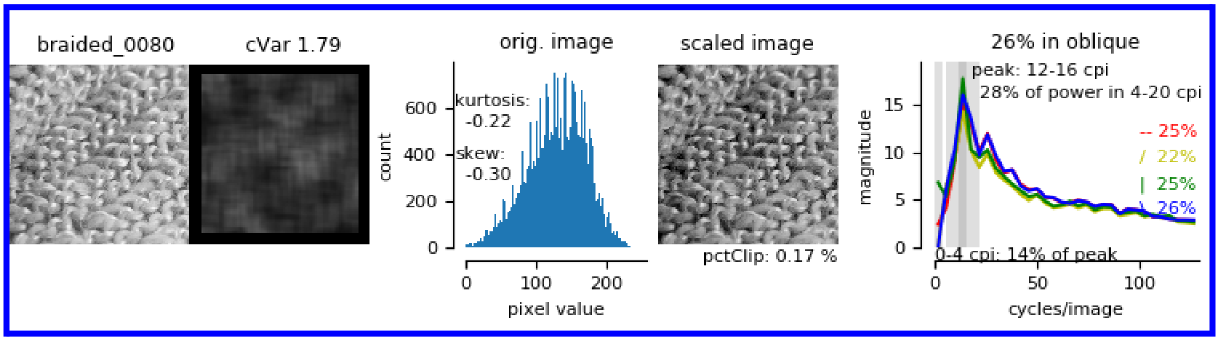
braided_0080
cVar 1.79
orig. image
kurtosis: -0.22
skew: -0.30
count
pixel value
scaled image
pctClip: 0.17 %
26% in oblique
peak: 12-16 cpi
28% of power in 4-20 cpi
-- 25%
/ 22%
| 25%
\ 26%
0-4 cpi: 14% of peak
magnitude
cycles/image

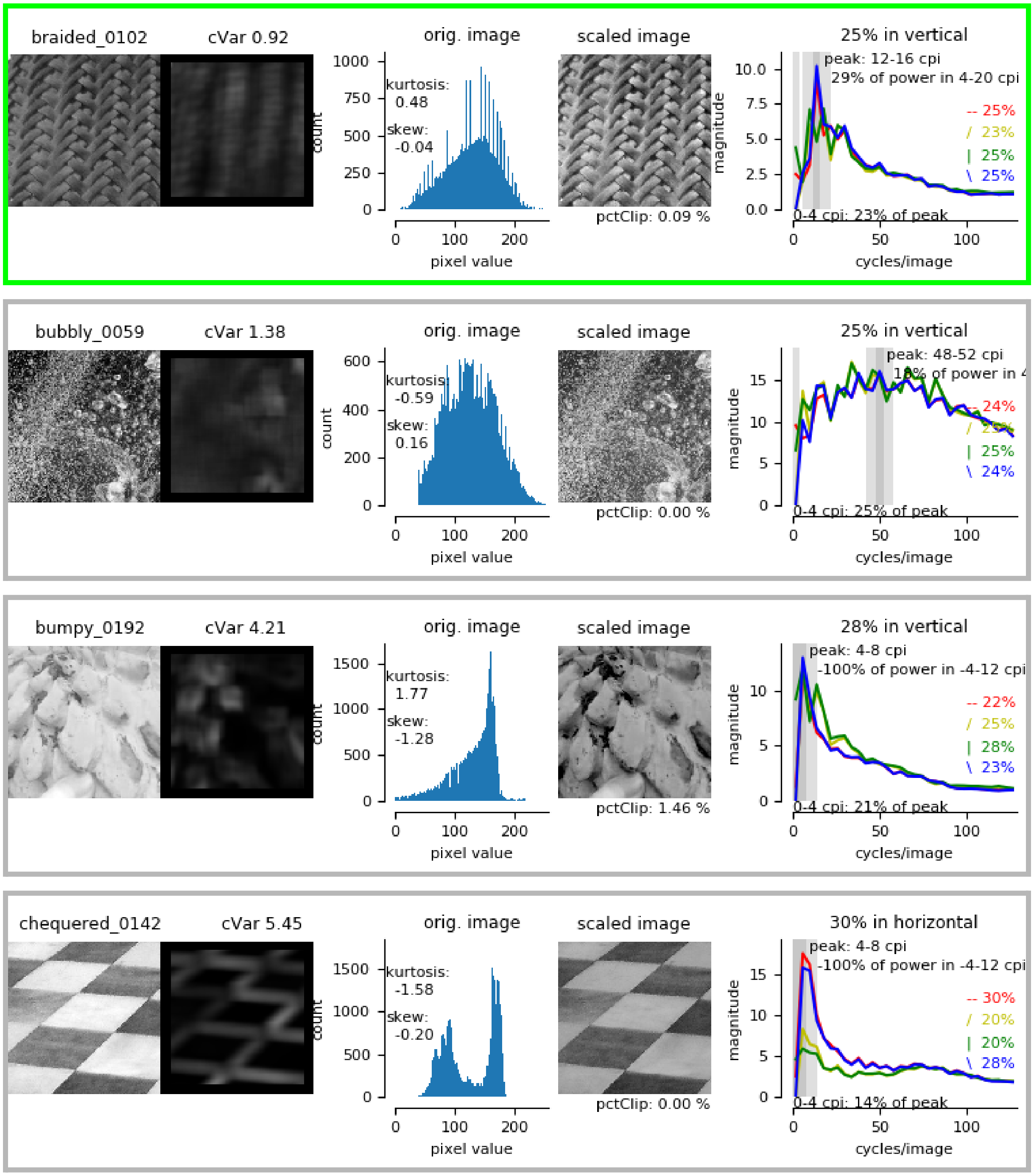

braided_0102
cVar 0.92
orig. image
kurtosis: 0.48
skew: -0.04
count
pixel value
scaled image
pctClip: 0.09 %
25% in vertical
peak: 12-16 cpi
29% of power in 4-20 cpi
-- 25%
/ 23%
| 25%
\ 25%
0-4 cpi: 23% of peak
magnitude
cycles/image
bubbly_0059
cVar 1.38
orig. image
kurtosis: -0.59
skew: 0.16
count
pixel value
scaled image
pctClip: 0.00 %
25% in vertical
peak: 48-52 cpi
-- 24%
| 25%
\ 24%
0-4 cpi: 25% of peak
magnitude
cycles/image
bumpy_0192
cVar 4.21
orig. image
kurtosis: 1.77
skew: -1.28
count
pixel value
scaled image
pctClip: 1.46 %
28% in vertical
peak: 4-8 cpi
-100% of power in -4-12 cpi
-- 22%
/ 25%
| 28%
\ 23%
0-4 cpi: 21% of peak
magnitude
cycles/image
chequered_0142
cVar 5.45
orig. image
kurtosis: -1.58
skew: -0.20
count
pixel value
scaled image
pctClip: 0.00 %
30% in horizontal
peak: 4-8 cpi
-100% of power in -4-12 cpi
-- 30%
/ 20%
| 20%
\ 28%
0-4 cpi: 14% of peak
magnitude
cycles/image

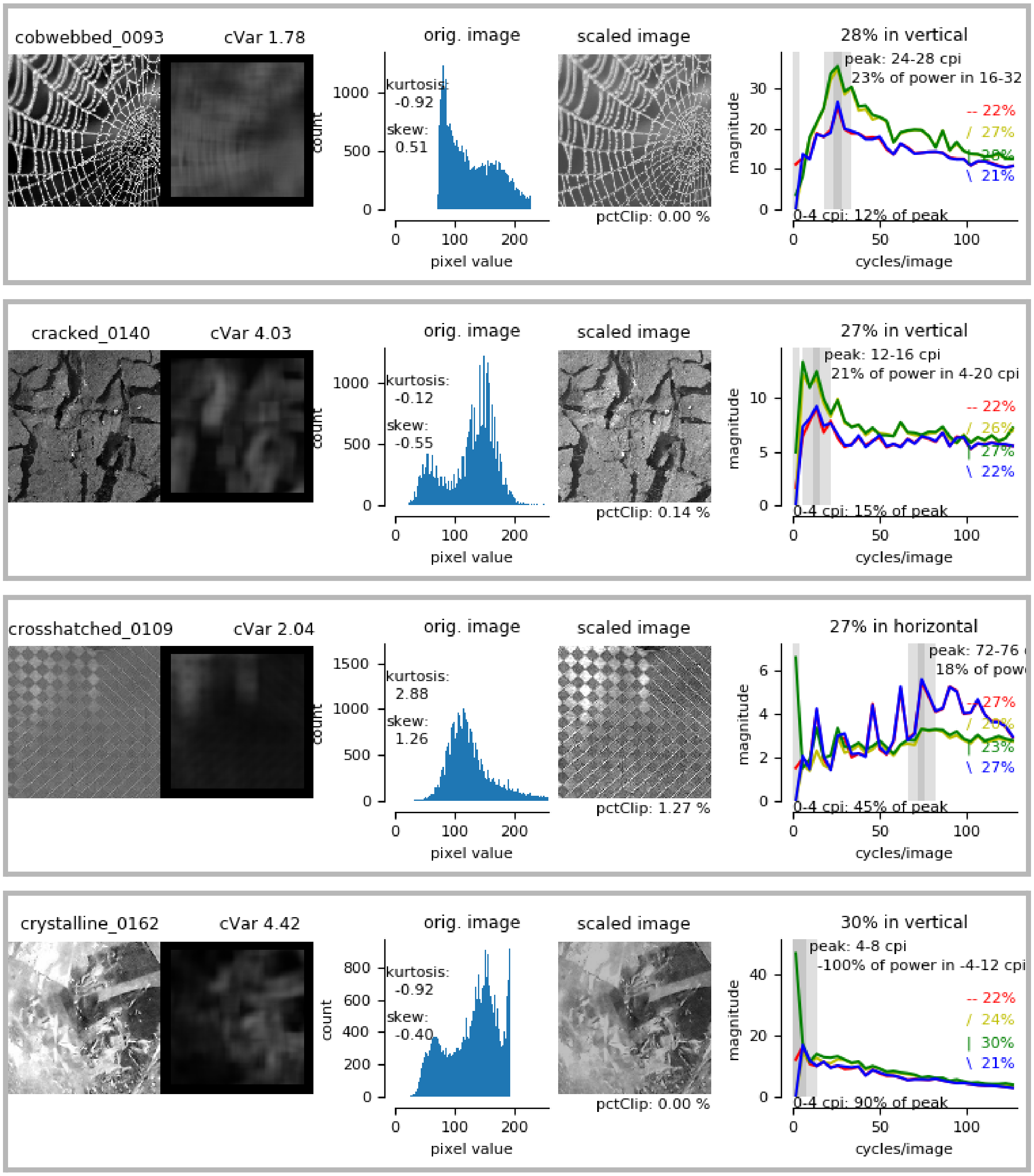
cobwebbed_0093
cVar 1.78
orig. image
kurtosis:
-0.92
skew:
0.51
count
pixel value
scaled image
pctClip: 0.00 %
28% in vertical
peak: 24-28 cpi
23% of power in 16-32
-- 22%
/ 27%
| 28%
\ 21%
0-4 cpi: 12% of peak
magnitude
cycles/image
cracked_0140
cVar 4.03
orig. image
kurtosis:
-0.12
skew:
-0.55
count
pixel value
scaled image
pctClip: 0.14 %
27% in vertical
peak: 12-16 cpi
21% of power in 4-20 cpi
-- 22%
/ 26%
| 27%
\ 22%
0-4 cpi: 15% of peak
magnitude
cycles/image
crosshatched_0109
cVar 2.04
orig. image
kurtosis:
2.88
skew:
1.26
count
pixel value
scaled image
pctClip: 1.27 %
27% in horizontal
peak: 72-76
18% of pow
-- 27%
/ 23%
| 23%
\ 27%
0-4 cpi: 45% of peak
magnitude
cycles/image
crystalline_0162
cVar 4.42
orig. image
kurtosis:
-0.92
skew:
-0.40
count
pixel value
scaled image
pctClip: 0.00 %
30% in vertical
peak: 4-8 cpi
-100% of power in -4-12 cpi
-- 22%
/ 24%
| 30%
\ 21%
0-4 cpi: 90% of peak
magnitude
cycles/image

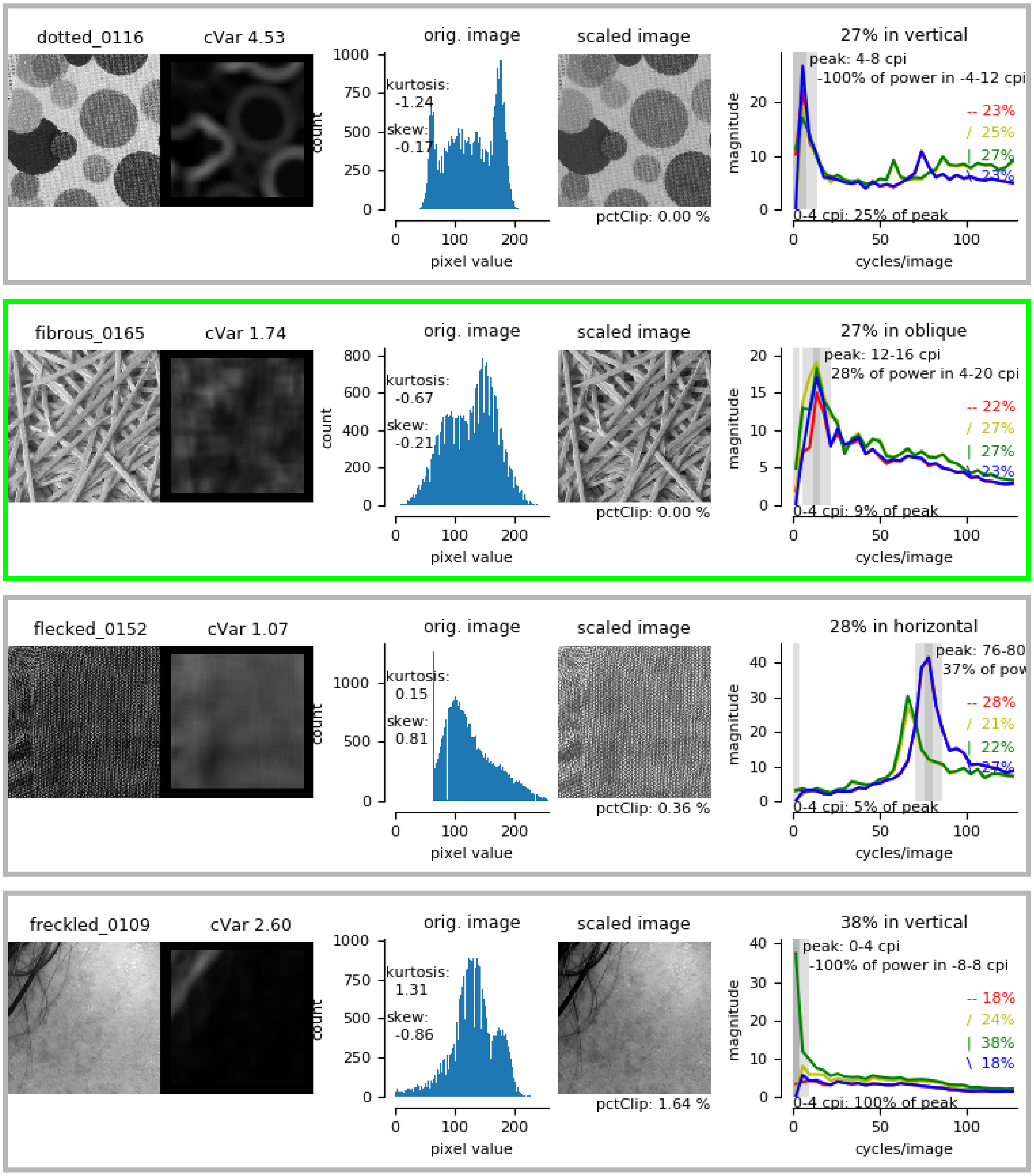
dotted_0116
cVar 4.53
orig. image
kurtosis: -1.24
skew: -0.17
count
pixel value
scaled image
pctClip: 0.00 %
27% in vertical
peak: 4-8 cpi
-100% of power in -4-12 cpi
-- 23%
/ 25%
| 27%
\ 23%
0-4 cpi: 25% of peak
magnitude
cycles/image
fibrous_0165
cVar 1.74
orig. image
kurtosis: -0.67
skew: -0.21
count
pixel value
scaled image
pctClip: 0.00 %
27% in oblique
peak: 12-16 cpi
28% of power in 4-20 cpi
-- 22%
/ 27%
| 27%
\ 23%
0-4 cpi: 9% of peak
magnitude
cycles/image
flecked_0152
cVar 1.07
orig. image
kurtosis: 0.15
skew: 0.81
count
pixel value
scaled image
pctClip: 0.36 %
28% in horizontal
peak: 76-80
37% of pow
-- 28%
/ 21%
| 22%
\ 27%
0-4 cpi: 5% of peak
magnitude
cycles/image
freckled_0109
cVar 2.60
orig. image
kurtosis: 1.31
skew: -0.86
count
pixel value
scaled image
pctClip: 1.64 %
38% in vertical
peak: 0-4 cpi
-100% of power in -8-8 cpi
-- 18%
/ 24%
| 38%
\ 18%
0-4 cpi: 100% of peak
magnitude
cycles/image

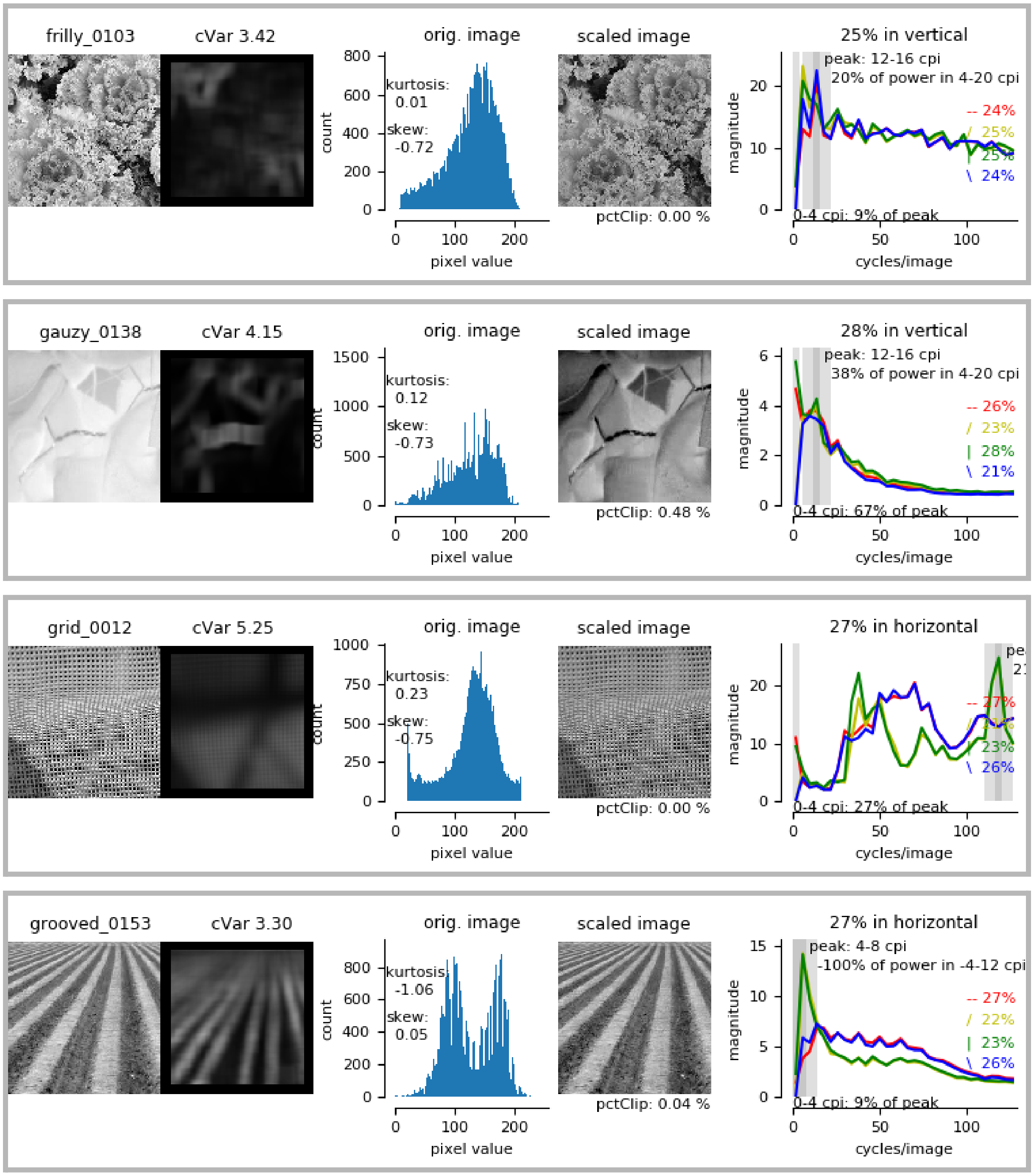
frilly_0103
cVar 3.42
orig. image
kurtosis:
0.01
skew:
-0.72
count
pixel value
scaled image
pctClip: 0.00 %
25% in vertical
peak: 12-16 cpi
20% of power in 4-20 cpi
-- 24%
/ 25%
\ 24%
magnitude
0-4 cpi: 9% of peak
cycles/image
gauzy_0138
cVar 4.15
orig. image
kurtosis:
0.12
skew:
-0.73
count
pixel value
scaled image
pctClip: 0.48 %
28% in vertical
peak: 12-16 cpi
38% of power in 4-20 cpi
-- 26%
/ 23%
| 28%
\ 21%
magnitude
0-4 cpi: 67% of peak
cycles/image
grid_0012
cVar 5.25
orig. image
kurtosis:
0.23
skew:
-0.75
count
pixel value
scaled image
pctClip: 0.00 %
27% in horizontal
-- 27%
| 23%
\ 26%
magnitude
0-4 cpi: 27% of peak
cycles/image
grooved_0153
cVar 3.30
orig. image
kurtosis:
-1.06
skew:
0.05
count
pixel value
scaled image
pctClip: 0.04 %
27% in horizontal
peak: 4-8 cpi
-100% of power in -4-12 cpi
-- 27%
/ 22%
| 23%
\ 26%
magnitude
0-4 cpi: 9% of peak
cycles/image

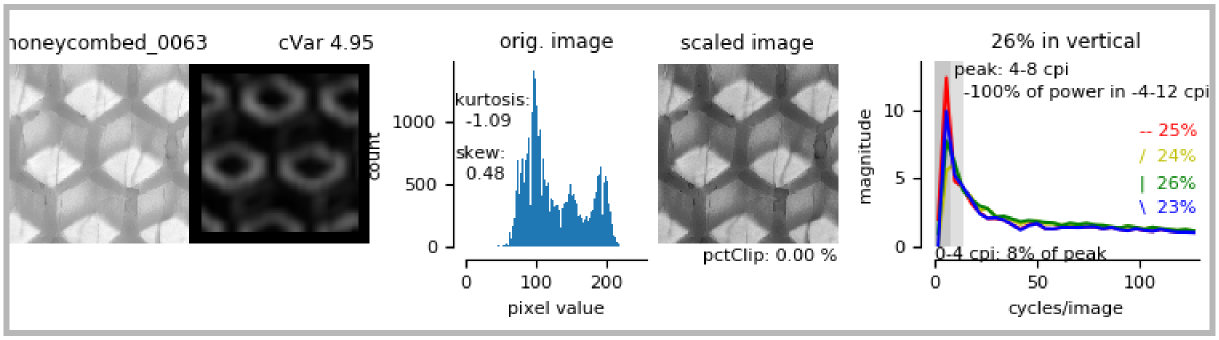
honeycombed_0063
cVar 4.95
orig. image
kurtosis:
-1.09
skew:
0.48
count
pixel value
scaled image
pctClip: 0.00 %
26% in vertical
peak: 4-8 cpi
-100% of power in -4-12 cpi
-- 25%
/ 24%
| 26%
\ 23%
0-4 cpi: 8% of peak
magnitude
cycles/image

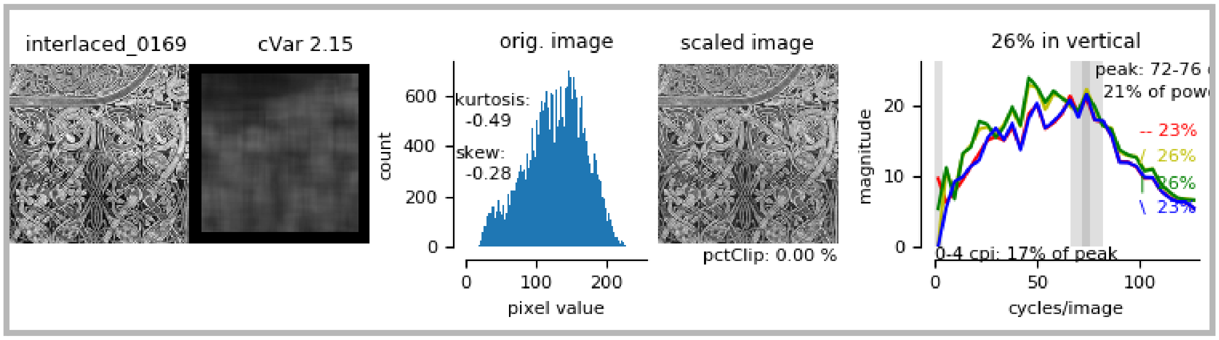
interlaced_0169
cVar 2.15
orig. image
kurtosis:
-0.49
skew:
-0.28
count
pixel value
scaled image
pctClip: 0.00 %
26% in vertical
peak: 72-76
21% of pow
-- 23%
/ 26%
| 26%
\ 23%
0-4 cpi: 17% of peak
magnitude
cycles/image

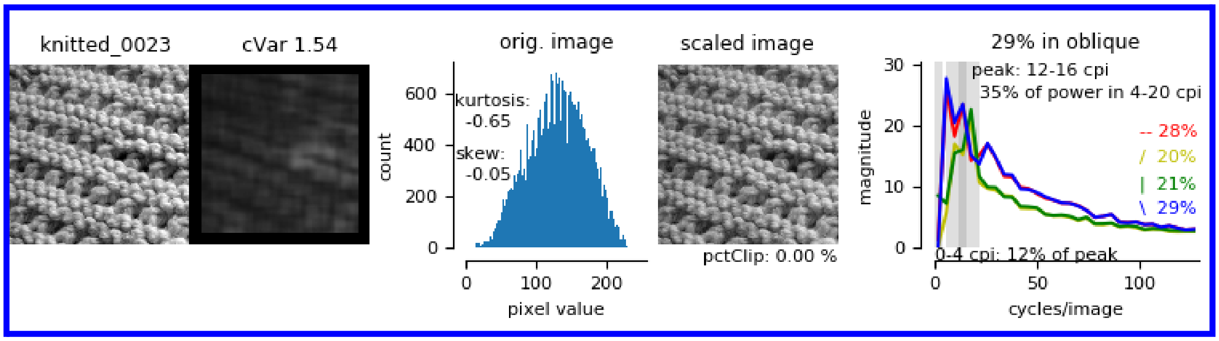
knitted_0023
cVar 1.54
orig. image
kurtosis:
-0.65
skew:
-0.05
count
pixel value
scaled image
pctClip: 0.00 %
29% in oblique
peak: 12-16 cpi
35% of power in 4-20 cpi
-- 28%
/ 20%
| 21%
\ 29%
0-4 cpi: 12% of peak
magnitude
cycles/image

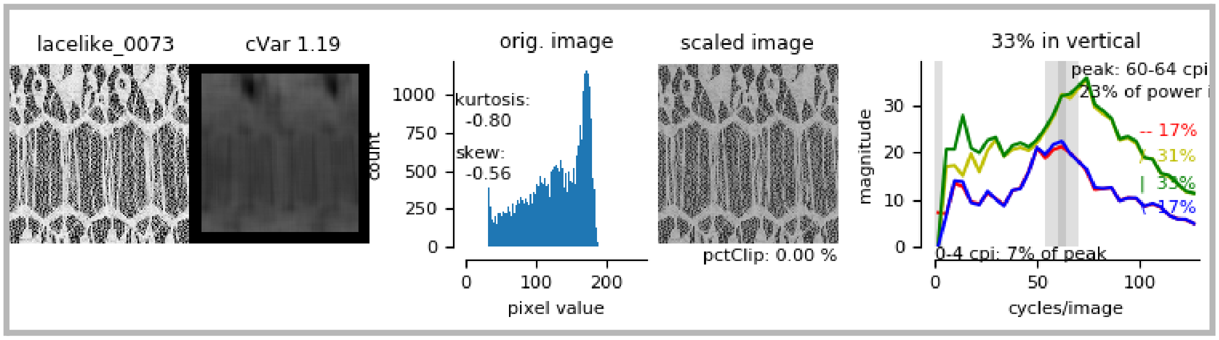
lacelike_0073
cVar 1.19
orig. image
kurtosis:
-0.80
skew:
-0.56
count
pixel value
scaled image
pctClip: 0.00 %
33% in vertical
peak: 60-64 cpi
23% of power
-- 17%
/ 31%
| 33%
\ 17%
0-4 cpi: 7% of peak
magnitude
cycles/image

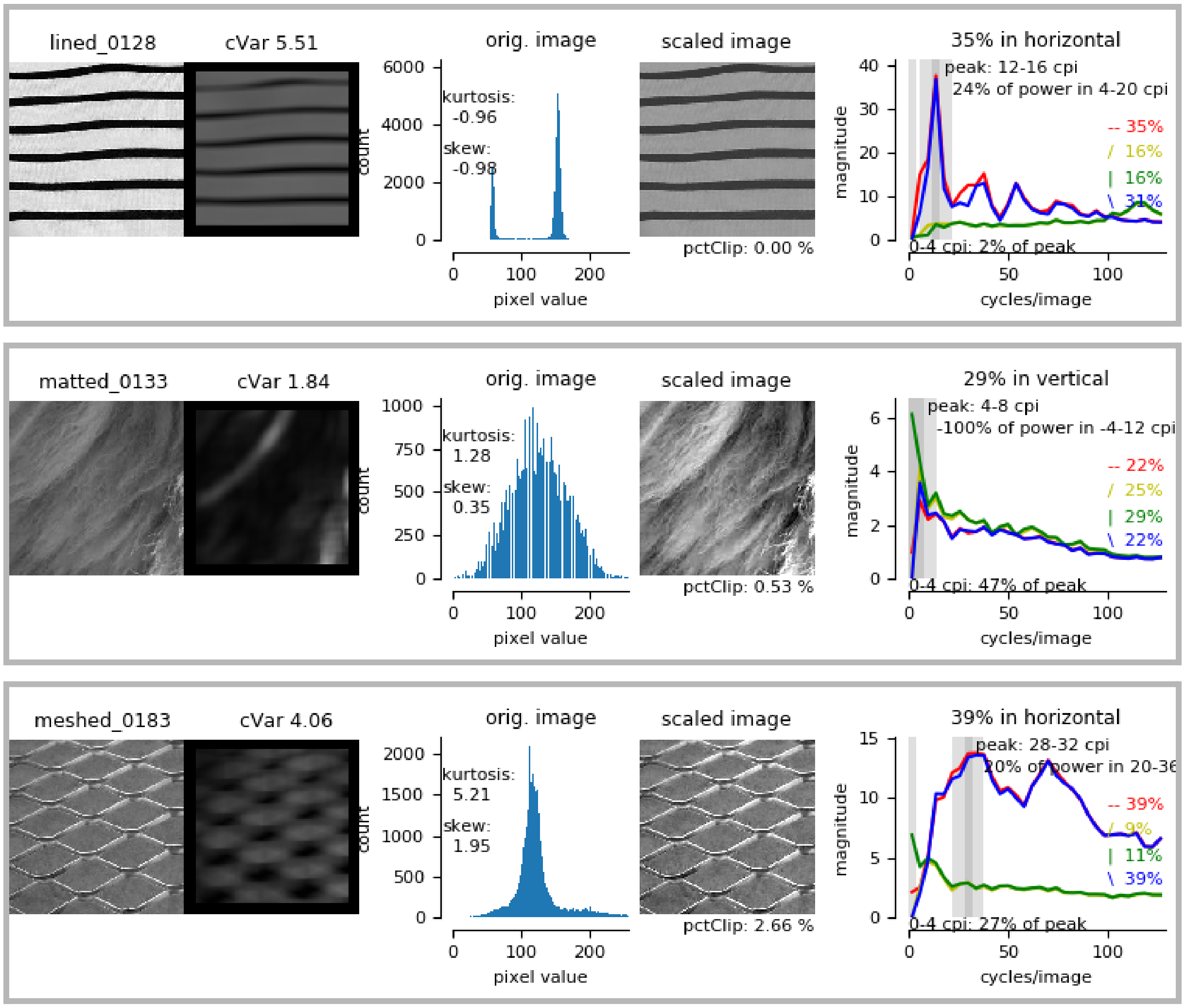

lined_0128
cVar 5.51
orig. image
kurtosis: -0.96
skew: -0.98
count
pixel value
scaled image
pctClip: 0.00 %
35% in horizontal
peak: 12-16 cpi
24% of power in 4-20 cpi
-- 35%
/ 16%
| 16%
\ 31%
0-4 cpi: 2% of peak
magnitude
cycles/image
matted_0133
cVar 1.84
orig. image
kurtosis: 1.28
skew: 0.35
count
pixel value
scaled image
pctClip: 0.53 %
29% in vertical
peak: 4-8 cpi
-100% of power in -4-12 cpi
-- 22%
/ 25%
| 29%
\ 22%
0-4 cpi: 47% of peak
magnitude
cycles/image
meshed_0183
cVar 4.06
orig. image
kurtosis: 5.21
skew: 1.95
count
pixel value
scaled image
pctClip: 2.66 %
39% in horizontal
peak: 28-32 cpi
20% of power in 20-36
-- 39%
/ 9%
| 11%
\ 39%
0-4 cpi: 27% of peak
magnitude
cycles/image

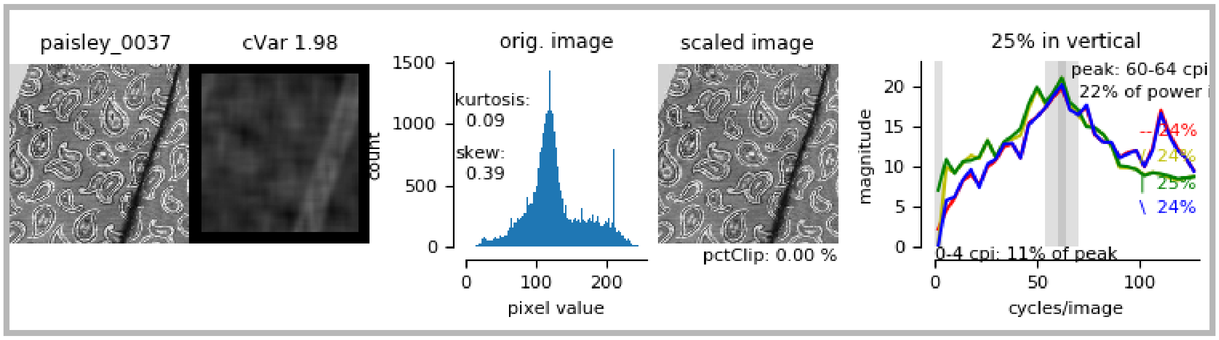
paisley_0037
cVar 1.98
orig. image
kurtosis: 0.09
skew: 0.39
count
pixel value
scaled image
pctClip: 0.00 %
25% in vertical
peak: 60-64 cpi
22% of power i
-- 24%
/ 24%
| 25%
\ 24%
0-4 cpi: 11% of peak
magnitude
cycles/image

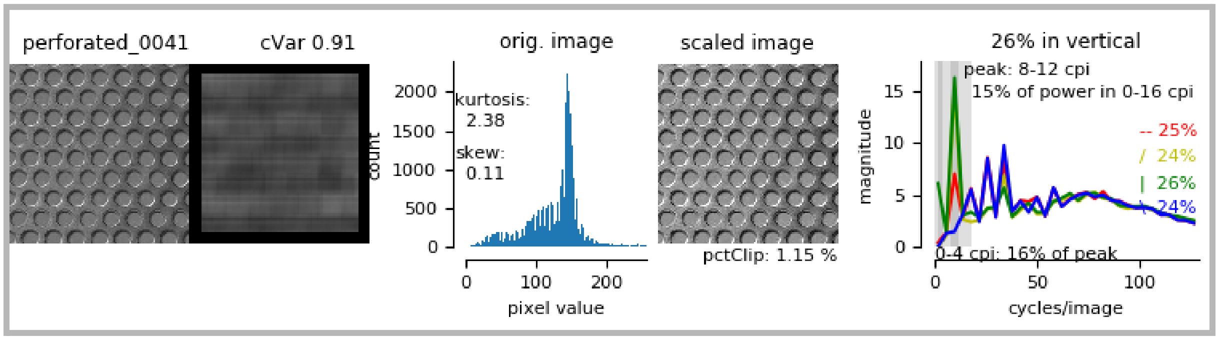
perforated_0041
cVar 0.91
orig. image
kurtosis: 2.38
skew: 0.11
count
pixel value
scaled image
pctClip: 1.15 %
26% in vertical
peak: 8-12 cpi
15% of power in 0-16 cpi
-- 25%
/ 24%
| 26%
\ 24%
0-4 cpi: 16% of peak
magnitude
cycles/image

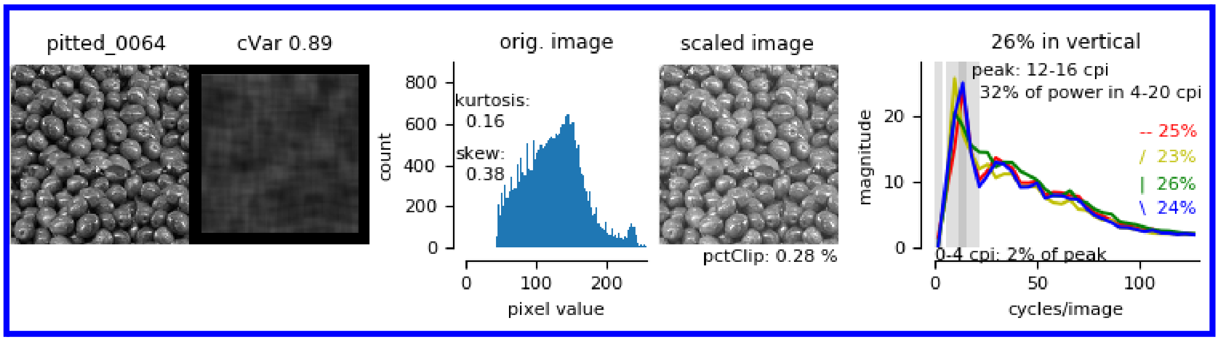
pitted_0064
cVar 0.89
orig. image
kurtosis: 0.16
skew: 0.38
count
pixel value
scaled image
pctClip: 0.28 %
26% in vertical
peak: 12-16 cpi
32% of power in 4-20 cpi
-- 25%
/ 23%
| 26%
\ 24%
0-4 cpi: 2% of peak
magnitude
cycles/image

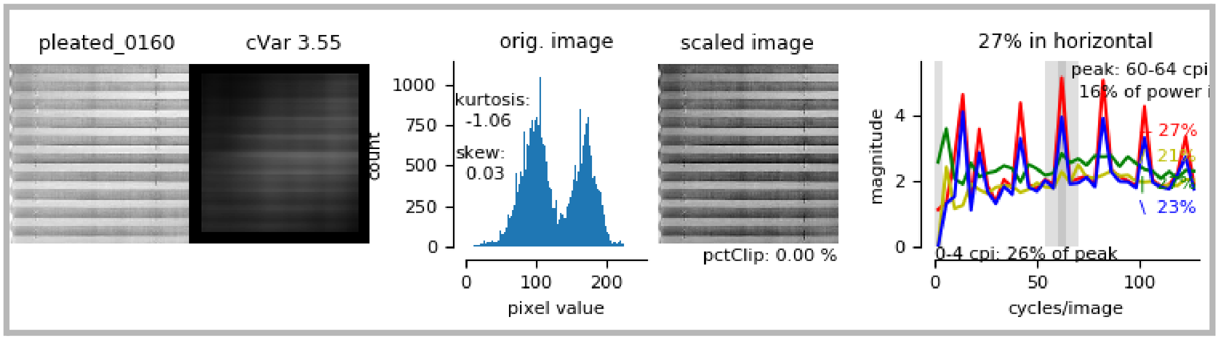
pleated_0160
cVar 3.55
orig. image
kurtosis: -1.06
skew: 0.03
count
pixel value
scaled image
pctClip: 0.00 %
27% in horizontal
peak: 60-64 cpi
16% of power i
-- 27%
\ 23%
0-4 cpi: 26% of peak
magnitude
cycles/image

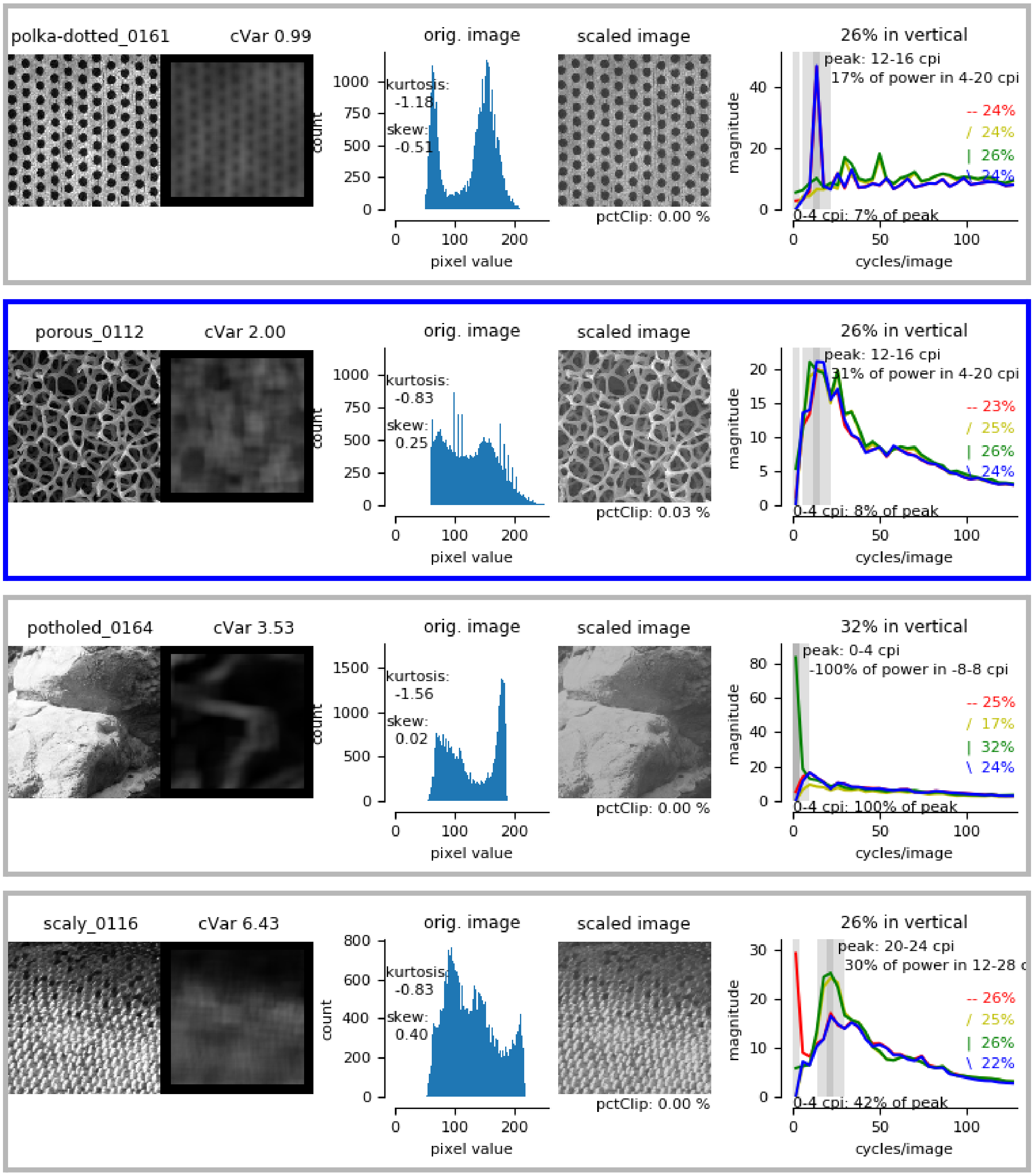

polka-dotted_0161
cVar 0.99
orig. image
kurtosis: -1.18
skew: -0.51
scaled image
pctClip: 0.00 %
26% in vertical
peak: 12-16 cpi
17% of power in 4-20 cpi
-- 24%
/ 24%
| 26%
\ 24%
0-4 cpi: 7% of peak
porous_0112
cVar 2.00
orig. image
kurtosis: -0.83
skew: 0.25
scaled image
pctClip: 0.03 %
26% in vertical
peak: 12-16 cpi
31% of power in 4-20 cpi
-- 23%
/ 25%
| 26%
\ 24%
0-4 cpi: 8% of peak
potholed_0164
cVar 3.53
orig. image
kurtosis: -1.56
skew: 0.02
scaled image
pctClip: 0.00 %
32% in vertical
peak: 0-4 cpi
-100% of power in -8-8 cpi
-- 25%
/ 17%
| 32%
\ 24%
0-4 cpi: 100% of peak
scaly_0116
cVar 6.43
orig. image
kurtosis: -0.83
skew: 0.40
scaled image
pctClip: 0.00 %
26% in vertical
peak: 20-24 cpi
30% of power in 12-28 c
-- 26%
/ 25%
| 26%
\ 22%
0-4 cpi: 42% of peak
count
pixel value
magnitude
cycles/image

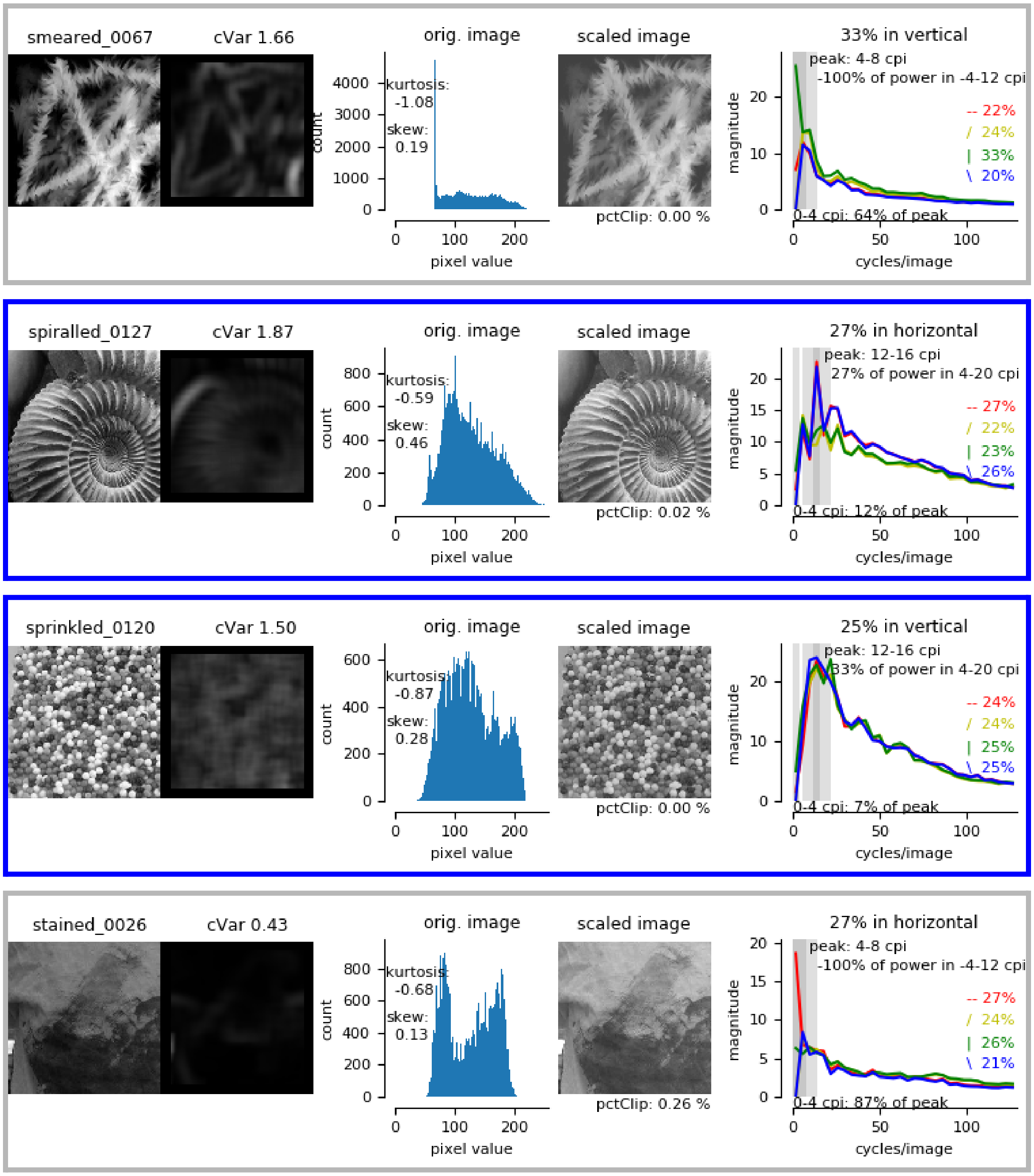
smeared_0067
cVar 1.66
orig. image
kurtosis:
-1.08
skew:
0.19
count
pixel value
scaled image
pctClip: 0.00 %
33% in vertical
peak: 4-8 cpi
-100% of power in -4-12 cpi
-- 22%
/ 24%
| 33%
\ 20%
0-4 cpi: 64% of peak
magnitude
cycles/image
spiralled_0127
cVar 1.87
orig. image
kurtosis:
-0.59
skew:
0.46
count
pixel value
scaled image
pctClip: 0.02 %
27% in horizontal
peak: 12-16 cpi
27% of power in 4-20 cpi
-- 27%
/ 22%
| 23%
\ 26%
0-4 cpi: 12% of peak
magnitude
cycles/image
sprinkled_0120
cVar 1.50
orig. image
kurtosis:
-0.87
skew:
0.28
count
pixel value
scaled image
pctClip: 0.00 %
25% in vertical
peak: 12-16 cpi
33% of power in 4-20 cpi
-- 24%
/ 24%
| 25%
\ 25%
0-4 cpi: 7% of peak
magnitude
cycles/image
stained_0026
cVar 0.43
orig. image
kurtosis:
-0.68
skew:
0.13
count
pixel value
scaled image
pctClip: 0.26 %
27% in horizontal
peak: 4-8 cpi
-100% of power in -4-12 cpi
-- 27%
/ 24%
| 26%
\ 21%
0-4 cpi: 87% of peak
magnitude
cycles/image

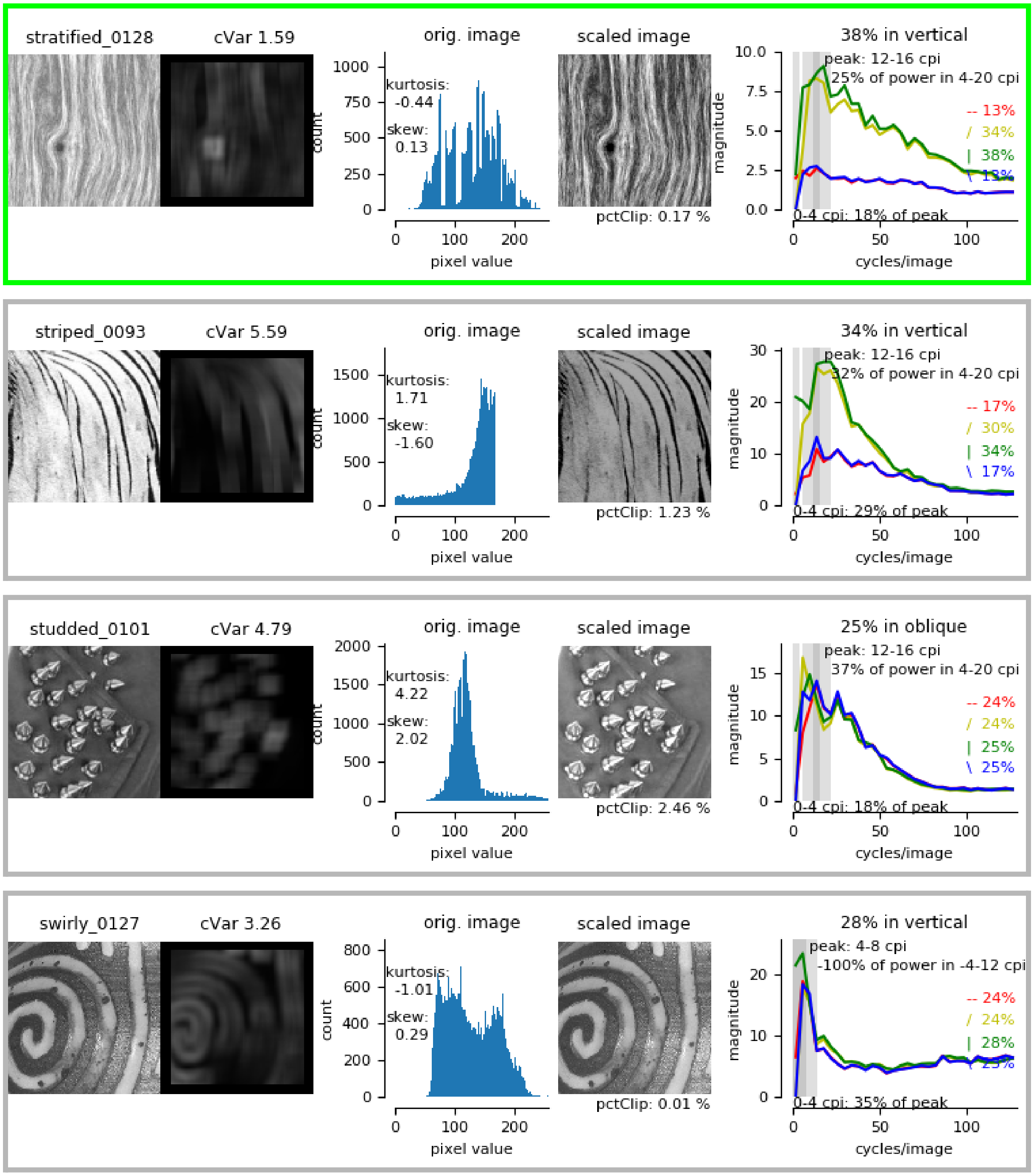

stratified_0128
cVar 1.59
orig. image
kurtosis: -0.44
skew: 0.13
count
pixel value
scaled image
pctClip: 0.17 %
38% in vertical
peak: 12-16 cpi
25% of power in 4-20 cpi
-- 13%
/ 34%
| 38%
\ 13%
0-4 cpi: 18% of peak
magnitude
cycles/image
striped_0093
cVar 5.59
orig. image
kurtosis: 1.71
skew: -1.60
count
pixel value
scaled image
pctClip: 1.23 %
34% in vertical
peak: 12-16 cpi
32% of power in 4-20 cpi
-- 17%
/ 30%
| 34%
\ 17%
0-4 cpi: 29% of peak
magnitude
cycles/image
studded_0101
cVar 4.79
orig. image
kurtosis: 4.22
skew: 2.02
count
pixel value
scaled image
pctClip: 2.46 %
25% in oblique
peak: 12-16 cpi
37% of power in 4-20 cpi
-- 24%
/ 24%
| 25%
\ 25%
0-4 cpi: 18% of peak
magnitude
cycles/image
swirly_0127
cVar 3.26
orig. image
kurtosis: -1.01
skew: 0.29
count
pixel value
scaled image
pctClip: 0.01 %
28% in vertical
peak: 4-8 cpi
-100% of power in -4-12 cpi
-- 24%
/ 24%
| 28%
0-4 cpi: 35% of peak
magnitude
cycles/image

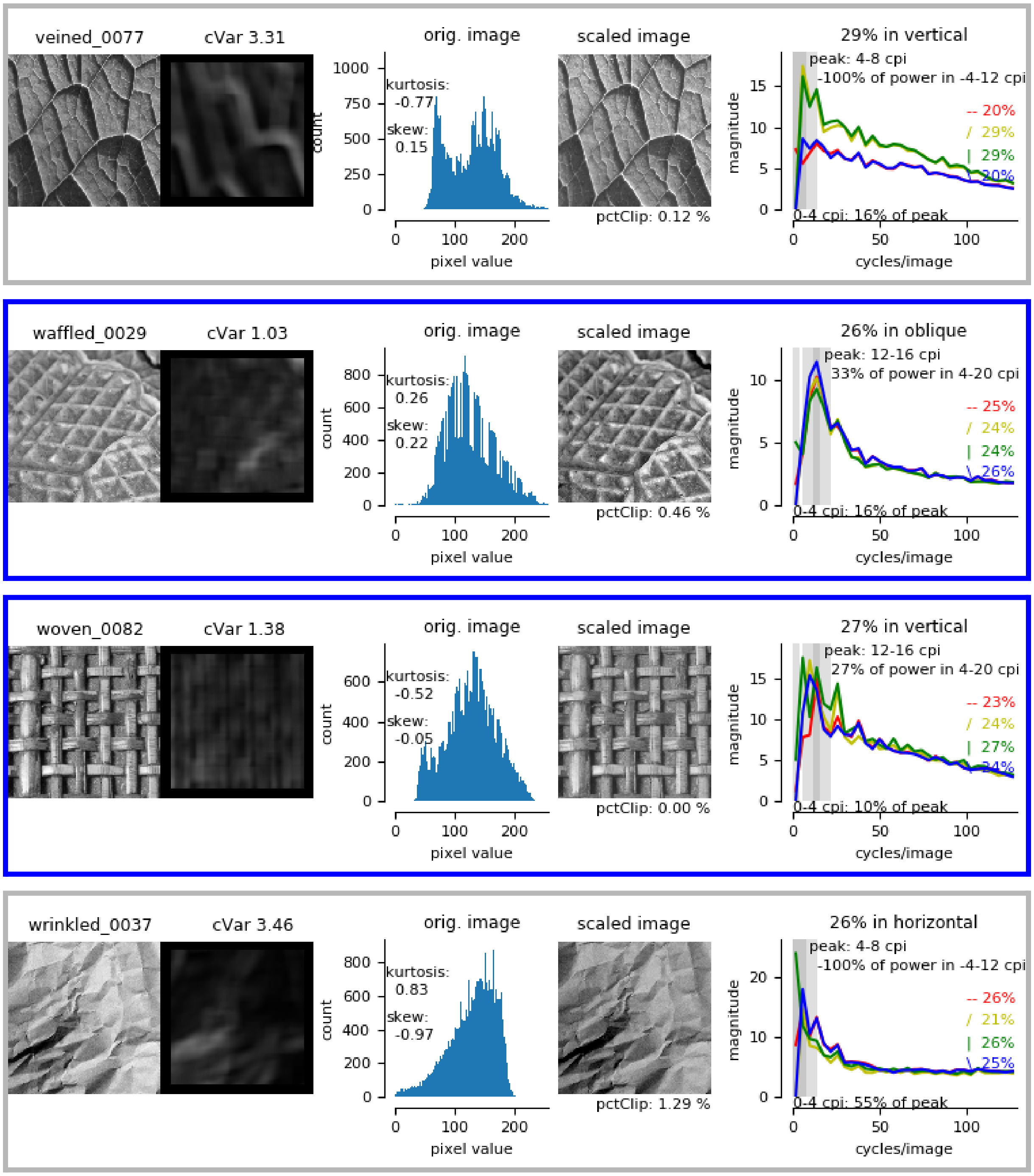
veined_0077
cVar 3.31
orig. image
kurtosis: -0.77
skew: 0.15
count
pixel value
scaled image
pctClip: 0.12 %
29% in vertical
peak: 4-8 cpi
-100% of power in -4-12 cpi
-- 20%
/ 29%
| 29%
\ 20%
0-4 cpi: 16% of peak
magnitude
cycles/image
waffled_0029
cVar 1.03
orig. image
kurtosis: 0.26
skew: 0.22
count
pixel value
scaled image
pctClip: 0.46 %
26% in oblique
peak: 12-16 cpi
33% of power in 4-20 cpi
-- 25%
/ 24%
| 24%
\ 26%
0-4 cpi: 16% of peak
magnitude
cycles/image
woven_0082
cVar 1.38
orig. image
kurtosis: -0.52
skew: -0.05
count
pixel value
scaled image
pctClip: 0.00 %
27% in vertical
peak: 12-16 cpi
27% of power in 4-20 cpi
-- 23%
/ 24%
| 27%
\ 24%
0-4 cpi: 10% of peak
magnitude
cycles/image
wrinkled_0037
cVar 3.46
orig. image
kurtosis: 0.83
skew: -0.97
count
pixel value
scaled image
pctClip: 1.29 %
26% in horizontal
peak: 4-8 cpi
-100% of power in -4-12 cpi
-- 26%
/ 21%
| 26%
\ 25%
0-4 cpi: 55% of peak
magnitude
cycles/image

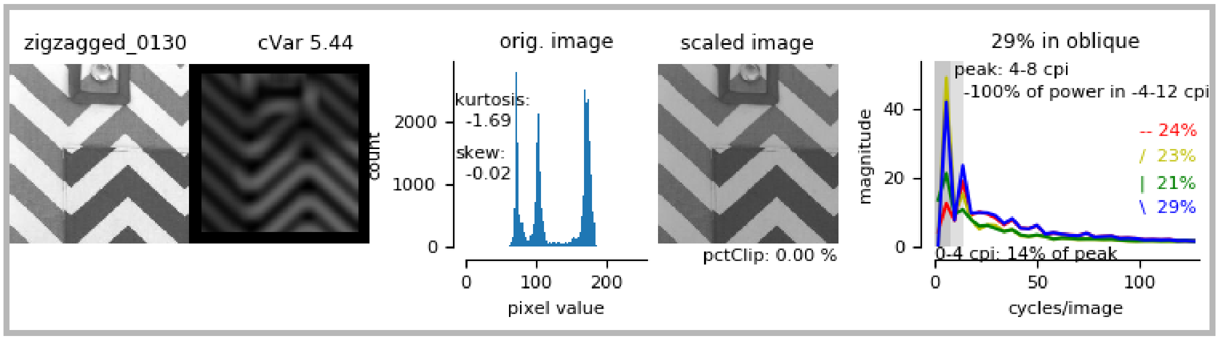


**Supplemental Figure 1. Graphical Summaries of Texture Image Statistics.** Gray borders indicate textures that were rejected in the first step. Blue borders indicate textures selected as candidates for consideration. Green borders indicate textures selected for the final experiment (green borders).

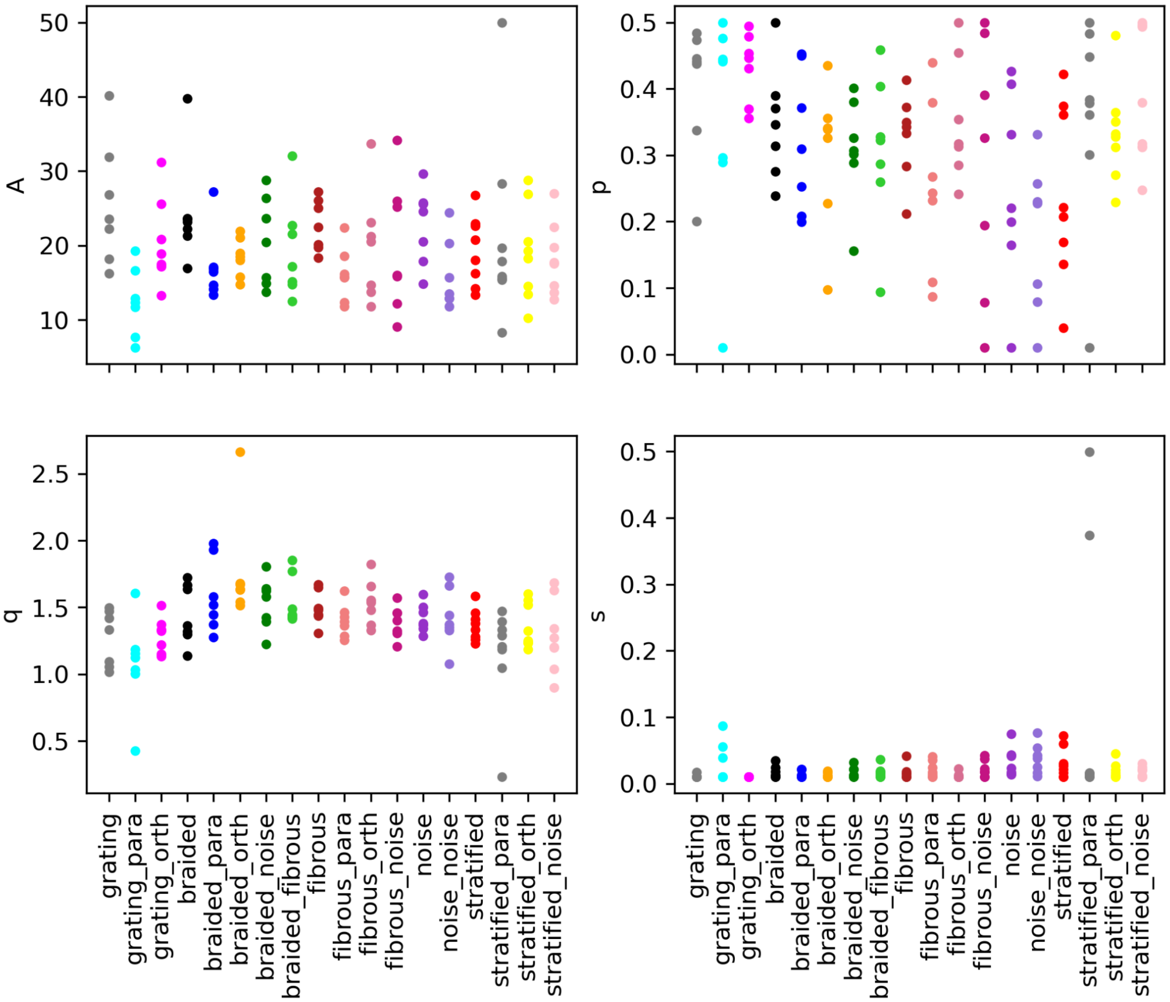


**Supplemental Figure 2. Naka-Rushton Parameters by Condition.** Dots represent each individuals' parameter value for each condition.
